# Parallel Activation of Src and Hif1α Increases Localized Glycolytic ATP Generation for Re-assembly of Endothelial Adherens Junctions

**DOI:** 10.1101/2022.11.11.516184

**Authors:** Li Wang, Priyanka Gajwani, Pallavi Chaturvedi, Zhigang Hong, Zijing Ye, Gregory J. Schwarz, Nicole M. Pohl-Avila, Anne-Marie Ray, Sarah Krantz, Peter T Toth, Deborah E. Leckband, Andrei Karginov, Jalees Rehman

## Abstract

Endothelial adherens junctions (AJs) are critical for the regulation of vascular barrier integrity and undergo dis-assembly during inflammatory injury, thus causing vascular leakiness. AJ re-assembly is thus necessary for restoration of the endothelial barrier following the initial injury. Here we examine the metabolic underpinnings that drive restoration of vascular integrity. In response to inflammatory stimuli, the glycolysis regulatory enzyme PFKFB3 is activated, resulting in a rapid and sustained increase of intracellular glycolytic ATP, especially in the proximity of AJs at the plasma membrane. We engineered a novel chemo-genetic construct (RapT) which allowed for precise temporal control of PFKFB3 recruitment to the plasma membrane. Activation of RapT by rapamycin during the barrier restoration phase increased regional ATP and accelerated AJ re-assembly. Mechanistically, we observed that PFKFB3 is activated through two modes. Src-mediated post-translational phosphorylation rapidly increases PFKFB3 activity. Using another chemo-genetic approach to temporally control Src activity, we found that Src activates PFKFB3 by binding to and phosphorylating it at residues Y175, Y334, and Y363. Tyrosine-phospho-deficient mutants of PFKFB3 at these residues block the glycolytic activation upon inflammatory stimuli. In parallel, elevated reactive oxygen species generated during inflammatory stimulation create pockets of regional hypoxia and allow for increased Hif1α-mediated transcription of PFKFB3, leading to sustained glycolytic activation. Moreover, inhibition of PFKFB3 delays AJ reassembly and restoration of vascular integrity both in vitro and in vivo. In conclusion, we show that while inflammatory activation acutely compromises the endothelial barrier, inflammatory signaling also concomitantly generates a metabolic milieu in anticipation of the subsequent re-assembly of AJs and restoration of the vascular barrier.

## Introduction

Vascular endothelial cells (ECs) control the transport of fluid, circulating proteins and immune cells into the tissues (Deanfield, Halcox et al. 2007). Adherens Junctions (AJs) are cell-cell adhesion complexes containing vascular endothelial cadherin (VE-cadherin), which are essential for the maintenance of the intact endothelial barrier, thus preventing uncontrolled fluid leak into the tissue (Yuksel, Ocalan et al. 2021). One devastating example of excessive fluid leak due to downregulation of AJs can be seen in the inflammatory lung injury during sepsis or severe SARS-CoV-2 infection. Inflammation-mediated AJs disruption leads to excessive neutrophil infiltration and protein-rich fluid leak into lung air space, resulting in edema and respiratory failure (Matthay, Zemans et al. 2019, Garde, Kenny et al. 2022, Zhang, Dutta et al. 2022). Thus, restoring the integrity of the endothelial barrier is a central factor determining the degree and extent of recovery (Zhang, Li et al. 2019, Evans, Iruela-Arispe et al. 2021). While it has been reported that inflammatory stimuli can upregulate glycolysis and induce inflammatory injury (Feng, Bowden et al. 2017, Zhang, Li et al. 2019, Xiao, Oldham et al. 2021), it is not known how metabolic shifts induced by inflammatory cues affect the restoration of vascular barrier integrity during the later resolution phase.

Vascular ECs mainly rely on glycolysis for rapid ATP production (De Bock, Georgiadou et al. 2013), thus avoiding consumption of oxygen designated for the tissues that are supplied by the vasculature and circumventing the generation of deleterious mitochondrial reactive oxygen species (ROS) that accompany oxidative phosphorylation (Vander Heiden, Cantley et al. 2009, Bolanos, Almeida et al. 2010, Metallo and Vander Heiden 2013). Several glycolytic enzymes such as hexokinase and 6-Phosphofructo-2-Kinase/Fructose-2,6-Biphosphatase 3 (PFKFB3) are key regulators of metabolic shift to meet the energy demand for cellular functions in the endothelium (De Bock, Georgiadou et al. 2013, Xu, An et al. 2014, Yu, Wilhelm et al. 2017). The product of PFKFB3, fructose-2,6-bisphosphate, is a potent activator of PFK-1, the key rate-limiting enzyme in the glycolysis pathway. PFKFB3 has been shown to promote filopodia/lamellipodia of endothelial tip cells and enhance blood vessel growth during angiogenesis, indicating an important role for this enzyme in EC function (De Bock, Georgiadou et al. 2013, Schoors, De Bock et al. 2014). However, the role of PFKFB3 in the regulation of EC barrier function during inflammation injury and subsequent recovery remain unclear.

PFKFB3 activity is tightly regulated by both transcriptional and post-translation mechanisms. One post-translational mechanism of PFKFB3 regulation is by serine/threonine phosphorylation (Yi, Ban et al. 2019). Recent studies point to a key role of the oncogenic tyrosine kinase Src in rewiring tumor metabolism towards glycolysis by activating glycolytic enzymes (Zhang, Wang et al. 2017, Ma, Zhang et al. 2020, Pelaz and Tabernero 2022). Src is also activated in ECs during inflammation (Alexander, Alexander et al. 2000), where it induces AJ permeability through direct phosphorylation of VE Cadherin (Sidibe and Imhof 2014). However, Src activity also has junction stabilizing effects, as pharmacological inhibition of Src results in reduced barrier intactness through an unknown mechanism (Kreutzman, Colom-Fernandez et al. 2017), and transient Src activation enhances endothelial barrier function (Klomp, Shaaya et al. 2019). This dual role of Src in destabilizing the endothelial barrier as well restoring the endothelial barrier integrity, together with its established role in activating glycolysis in tumors, suggests a possible role for Src in regulating endothelial metabolism and barrier integrity restoration via PFKFB3.

Here, we demonstrate that ECs respond to inflammatory stimuli by increasing glycolysis via PFKFB3 activation through two modes: a newly identified rapid post-translational mechanism of Src-mediated tyrosine phosphorylation, and a more sustained mechanism of Hif1α-mediated PFKFB3 transcription. Using RapR-Src, an engineered construct that allows for precise temporal regulation of Src, we show that Src binds, phosphorylates, and activates PFKFB3. Spatial control of PFKFB3 localization using a novel chemo-genetic engineered construct, RapT, shows that increased PFKFB3 activity at the membrane can accelerate restoration of the endothelial barrier. PFKFB3 inhibition during the resolution phase delayed the recovery of barrier function both in vivo and in vitro, thus highlighting the barrier restoration role for Src during the recovery period. We therefore posit that while inflammatory activation acutely compromises the EC barrier, inflammatory signaling concomitantly induces a metabolic shift towards increased glycolysis which promotes the restoration of the endothelial barrier and the resolution of tissue edema as well as inflammation.

## Results

To study endothelial glycolysis during inflammation, we first examined endothelial cell programming of glycolysis in response to inflammation. The pro-inflammatory cytokine TNFα is released by monocytes in response to inflammation and is a critical initiator of endothelial inflammatory signaling (Blaser, Dostert et al. 2016). We therefore investigated the effect of TNFα on the metabolic profile of human lung microvascular ECs (HLMVECs) by examining the ECAR (extracellular acidification rate, an indicator of glycolytic activity) and OCR (oxygen consumption rate, an indicator of mitochondrial respiration). We observed that TNFα increased the rate of glycolysis (**Figure 1a** and **Figure 1 supplementary Figure 1a**) but not that of mitochondrial respiration (**Figure 1 supplementary Figure 1c**). This glycolytic activation occurred as early as 15 minutes and was sustained for 6 hours, the duration of the experiment. Detailed ECAR analyses suggest that TNFα increases both glycolysis and glycolytic capacity but not the glycolytic reserve (**Figure 1 supplementary Figure 1b**).

**Figure 1.**
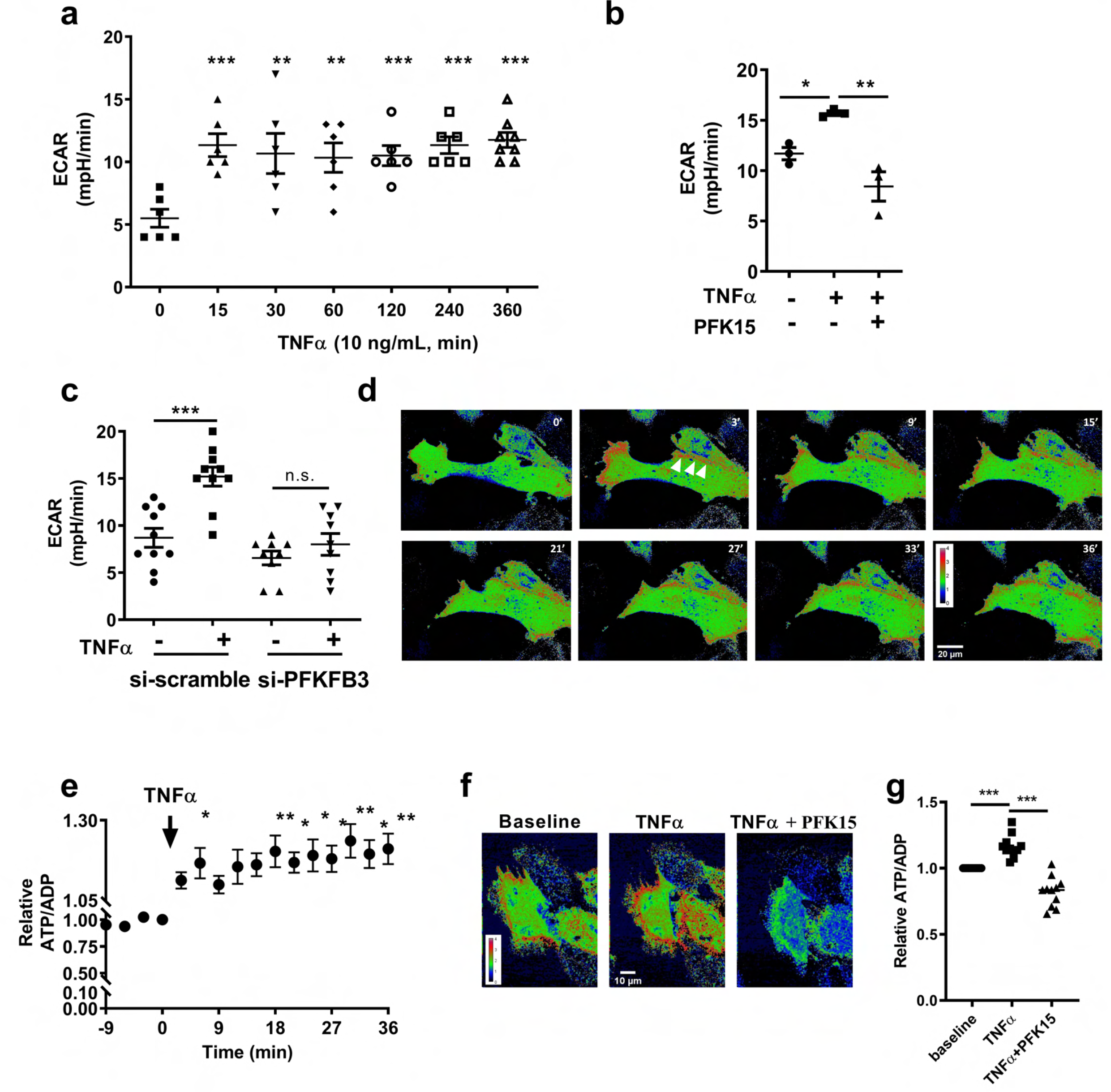
TNFα activates endothelial glycolysis and increases intracellular ATP via PFKFB3. (**a**)Time course of extracellular acidification rate (ECAR) measurements in HLMVECs showing that TNFα induces a rapid and persistent glycolytic activation. HLMVECs were treated with 10ng/ml TNFα, and ECAR was measured at the indicated time using Seahorse. (**b**) PFK15, a PFKFB3 specific inhibitor, prohibits TNFα-induced ECAR increases. HLMVECs were treated with TNFα, TNFα plus PFK15 (5 μM), or 0.1% DMSO as a vehicle control. The ECAR at 20 minutes after treatment was plotted. (**c**) TNFα-induced glycolytic activation is inhibited by PFKFB3-specific siRNAs in HLMVECs. HLMVECs were transfected with PFKFB3 siRNA. Scramble siRNA was used as a negative control. 48 hours post-transfection, cells were treated with TNFα. ECAR was measured 15 minutes after treatment. The successful knockdown of PFKFB3 is shown in Figure 1 supplementary figure 2. (**d**) Representative images of TNFα induced ATP/ADP changes at various time points, showing TNFα increases intracellular ATP/ADP spatially and temporally. The arrows indicate the regions of intracellular ATP/ADP increase. (**e**) Quantification showing TNFα increases intracellular ATP/ADP. The graph is calculated from movies of 10 cells in three-independent experiments. Statistical one-way ANOVA analysis shows statistically significant increases at 6, 18, 21, 24, 27, 30, 33, 36 minutes after TNFα stimulation. (**f**) Representative images showing that PFK15 inhibits TNFα-induced intracellular ATP/ADP increase. HLMVECs expressing Perceval HR were treated with TNFα (10 ng/mL) for 42 min, then followed by PFK15 (5 μM) treatment. Images of TNFα at 9 minutes and PFK15 after 15 minutes are shown. (**g**) Quantification showing PFK15 inhibits TNFα-induced intracellular ATP/ADP increase (n=11). Source data for (a) (b), (c), (e), and (g) are available in Figure 1 Source Data 1.

We next examined whether the glycolysis regulatory enzyme PFKFB3 was required for this EC metabolic response to TNFα. PFK15, a derivative of 3PO, another selective PFKFB3 inhibitor, is more potent and has better pharmacokinetic properties (Clem, O’Neal et al. 2013), was used to inhibit PFKFB3. Both pharmacological PFKFB3 inhibition by PFK15 and downregulation by si-PFKFB3 (**Figure 1 supplementary Figure 2**) attenuated TNFα-mediated glycolytic activation (**Figure 1b, c**). The fluorescent biosensor PercevalHR ratiometrically measures the intracellular ATP/ADP ratio (Tantama, Martinez-Francois et al. 2013), and was used to explore spatial and temporal intracellular ATP changes upon inflammatory stimulation. Expression of Perceval HR in HLMVECs allows for the monitoring of intracellular ATP levels as the addition of extracellular glucose increased intracellular ATP/ADP, and this increase in ATP production was diminished by PFKFB3 inhibition (**Figure 1 supplementary Figure 3**), confirming glycolytic ATP is the main energy resource in EC (De Bock, Georgiadou et al. 2013). Administration of TNFα rapidly induced a sustained increase in intracellular ATP (**Figure 1d, e**), which was abrogated by PFKFB3 inhibition (**Figure 1f, g**). These results signify that TNFα-induced upregulation of glycolytic ATP production, and this increase is mediated by PFKFB3.

The re-assembly of disrupted AJs is required for the restoration of homeostasis during the resolution of inflammation (Giannotta, Trani et al. 2013, Yan, Wang et al. 2016). This process included the recycling of AJ proteins, including VE-cadherin, back to the plasma membrane and the formation of junctions, which is an energy-consuming process. We therefore investigated the role of PFKFB3 in regulating AJ re-assembly by studying endothelial barrier permeability and membrane localization of VE-cadherin. During homeostasis, inhibition of PFKFB3 increased EC permeability in a dose-dependent manner (**Figure 2a, b**). VE-cadherin undergoes continuous turnover in cycles of endocytosis, sorting and recycling back to the plasma membrane (Giannotta, Trani et al. 2013). Thus, glycolytic inhibition may interfere with the energy-dependent VE-cadherin recycling, modulating EC barrier function. Visualization of membrane-localized VE-cadherin demonstrated a dose-dependent disruption of VE-cadherin at the AJs by PFKFB3 inhibition (**Figure 2c, d**). To test the functionality of adherens junctions in PFKFB3-inhibited cells, we measured the tension on VE-cadherin proteins within AJs, by employing a fluorescence-based, VE-cadherin tension sensor (VE-cad-TS) (**Figure 2e**), as loss of tension at cadherin complexes is associated with increased permeability (Gomez, McLachlan et al. 2011, Conway, Breckenridge et al. 2013, Buckley, Tan et al. 2014). Treatment of cell pairs with PFK15 for 40 minutes significantly reduced the tension on VE-cadherin in AJs between cell pairs (**Figure 2f, g**). Thus, loss of PFKFB3 activity results in reduced tension on VE-cadherin complexes, as well as in the reduced level of membrane VE Cadherin, leading to increased permeability. Here we showed that PFKFB3-mediated glycolysis is required for stabilizing AJs.

To study the role of glycolysis in restoring disrupted barriers, we next induced non-inflammatory injury by the addition of the extracellular calcium (Ca^2+^_ex_) chelator EDTA, as Ca^2+^_ex_ is necessary to maintain extracellular VE-cadherin interactions at the junction (Le, Yap et al. 1999, Yan, Wang et al. 2016). EDTA triggered membrane VE-cadherin disruption (**Figure 3a, b**), resulting in decreased trans-endothelial electrical resistance (TEER) (**Figure 3c, d**), whereas addition of Ca^2+^ restored AJs and endothelial barrier integrity (**Figure 3a-d)**. Concomitant inhibition of PFKFB3, however, prevented the restoration of EC barrier function (**Figure 3a-d)**. Additionally, trans-endothelial resistance returned to baseline once the selective PFKFB3 inhibitor PFK15 was washed out, demonstrating that this inhibition was reversible (**Figure 3c, e**). We also observed a similar effect with 2-DG, a global glycolysis inhibitor which acts upstream of PFKFB3 by reducing glucose entry into the glycolysis pathway (**Figure 3 supplementary Figure 1**). The inflammatory mediator Thrombin induces endothelial permeability in a rapid and reversible way (Rabiet, Plantier et al. 1996) and was thus used to study the effect of PFKFB3 overexpression in EC permeability. PFKFB3 overexpression in HLMVECs (**Figure 3 supplementary Figure 2a**) reduced thrombin-induced permeability and enhanced the recovery (**Figure 3 supplementary Figure 2b, c**). These results highlight the critical role of glycolysis in re-establishing EC barrier function following AJs disruption.

We next examined the effect of PFKFB3 inhibition on the restoration of EC barrier in an in vivo model of inflammatory endothelial injury and barrier disruption. As genetic deletion of PFKFB3 does not allow for targeted temporal inhibition of PFKFB3 during the barrier restoration phase, we used PFK15 to inhibit PFKFB3 only during resolution of inflammation in an endotoxemia model of lung endothelial injury. Systemic LPS injection in C57/BL6 mice induced peak vascular leak (as assessed by lung trans-vascular leak of Evans’ blue conjugated albumin, EBA) into lungs at 24 hours post challenge and the barrier integrity returned to baseline levels by 72 hours (**Figure 2h**). PFK15 was thus administered at the peak injury time point of 24 hours post LPS challenge to inhibit PFKFB3 mediated glycolysis exclusively during the injury resolution and barrier restoration phase. We observed markedly higher vascular leak (increased EBA) in the lungs of PFK15-treated mice at both 48 hours and 72 hours post LPS-administration, suggesting that PFKFB3 inhibition prevented the recovery of endothelial barrier function (**Figure 2h**). PFKFB3 inhibition during the resolution phase also impaired the survival of LPS-treated mice (**Figure 2i**), suggesting a requisite role for PFKFB3-mediated glycolysis in the vascular barrier repair post-injury.

To better understand how PFKFB3 responds to inflammation, we next investigated the mechanism by which TNFα activates PFKFB3. TNFα significantly increased PFKFB3 mRNA in HLMVECs starting at 1 hour (**Figure 4 supplementary Figure 1**) and PFKFB3 protein at 3 hours (**Figure 4a**). Analysis of the human PFKFB3 promoter region identified four putative hypoxia response elements. In addition, we observed that Hif1α protein level was significantly increased starting at 30 minutes after TNFα treatment (**Figure 4b**), preceding the observed PFKFB3 mRNA increase. Targeted depletion of Hif1α by siRNA (**Figure 4 supplementary Figure 2**) inhibited TNFα-mediated upregulation of PFKFB3 mRNA (**Figure 4c**), indicating that the observed increase in PFKFB3 mRNA is driven by the transcription factor Hif1α. Moreover, Hif1α activation was sufficient to drive PFKFB3 transcription even in the absence of TNFα as the hypoxia mimic DMOG increased PFKFB3 mRNA levels (**Figure 4c**). This is consistent with prior reports that Hif1α can transcriptionally upregulate PFKFB3 in other cell types (Minchenko, Leshchinsky et al. 2002). To explore how TNFα increases Hif1α levels, we took advantage of a ratiometric cell permeable oxygen sensor, MM2 (Dussmann, Perez-Alvarez et al. 2017). TNFα stimulation created subcellular pockets of reduced intracellular oxygen levels, comparable to what is seen when exposing cells to low exogenous oxygen conditions (**Figure 4d**). These results suggest that TNFα induces intracellular hypoxia, likely by the known consumption of O_2_ to generate superoxide radicals via NADPH oxidases (Smith, Waypa et al. 2017), thus inducing intracellular pockets of hypoxia which activate Hif1α-dependent PFKFB3 transcription.

**Figure 2.**
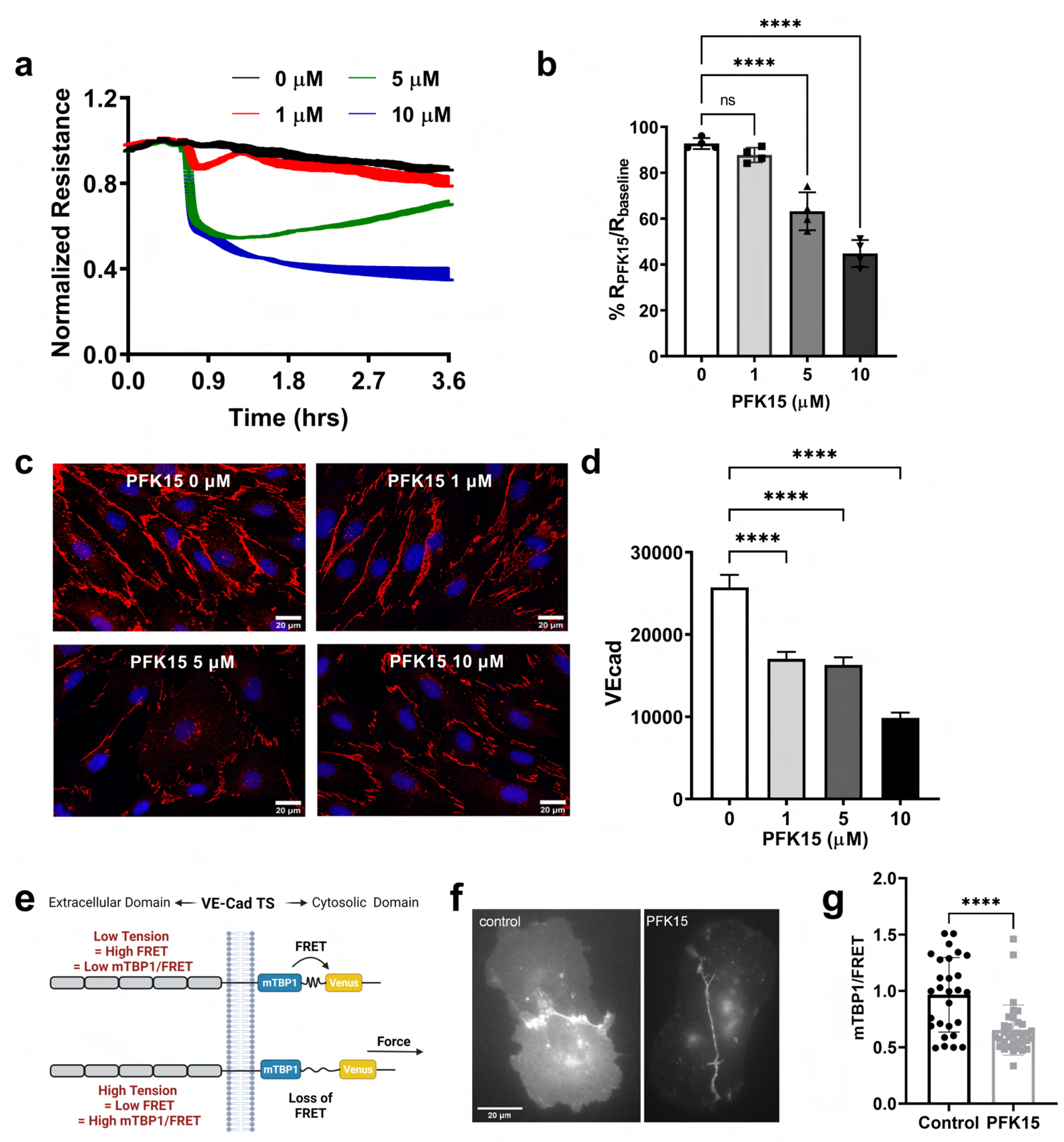
PFKFB3 stabilizes the endothelial barrier. (**a-b**) TEER measurement showing PFK15 dose-dependently inhibits EC barrier. The resistance of HLMVECs in HBSS was recorded using an ECIS instrument. Once stabilized, cells were treated with various doses of PFK15. Representative TEER measurement is shown in a. The resistance was normalized to the baseline, and the resistance affected by PFK15 was calculated by dividing resistance at two hours after PFK15 treatment (R_PFK15_) with baseline before treatment (R_baseline_) and is shown in b. (**c-d**) VE-cadherin staining of HLMVECs showing that PFK15 dose-dependently diminishes VE-cadherin at the plasma membrane. 45 minutes after PFK15 treatment, cells were fixed and stained with VE-cadherin antibody for detection of membrane VE-cadherin. Representative cell images (Red: VE-cadherin, Blue: DAPI) are shown in c and quantification of total membrane VE-cadherin fluorescence per cell are shown in d. (**e**) Schematic of VE-Cad-TS measuring junction tension. Loss of the FRET indicates increased junction tension (**f**) images of the Venus channel, showing junctional localization of VE-Cad-TS between cell pairs. (**g**) PFK15 reduces the junction tension. Cells were treated with 10μM PFK15 for 40 minutes, and junction tension was quantified (mTBP1/FRET, n=34) and compared to untreated cells (n = 29). Source data for (a) (b), (d), and (g) are available in Figure 2 Source Data 1.

**Figure 3.**
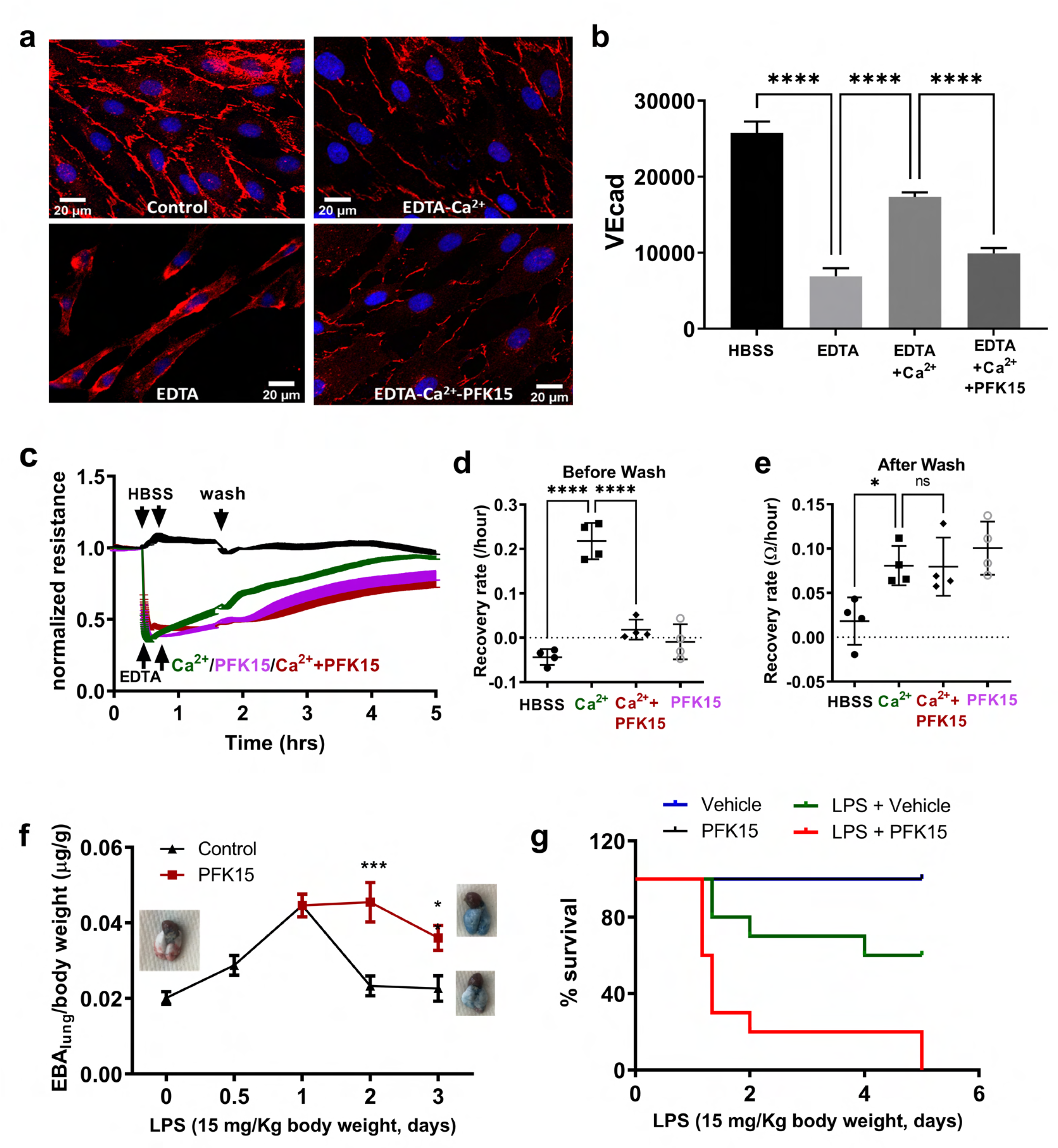
PFKFB3 enhances the recovery of barrier function after injury. (**a-b**) Immunofluorescent staining of VE-cadherin showed that PFK15 delays reannealing of AJs. HLMVECs were treated with EDTA (1mM) for 15 minutes to disrupt AJs, followed by adding Ca^2+^ (1mM) and PFK15 (10 μM). 30 minutes later, cells were fixed and stained for VE-cadherin. Representative images are shown in a and quantification of total membrane VE-cadherin immunofluorescence per cell are shown in b. (**c-e**) TEER measurement showing PFK15 inhibits the restoration of EC permeability. Cells were treated as in a-b, with an additional wash step during the recovery to remove PFK15 at e. Cells were monitored until resistance was recovered. Representative TEER recording is shown in c. The changes in relative resistance over a period (20 minutes) were calculated and are shown in d (before wash) and in e (after wash). (**f**) Time course of EBA permeability assay showing PFK15 delays recovery of lung endothelial barrier function upon LPS challenge. Intraperitoneal (IP) injection with LPS (15 mg/Kg) to C57/bl6 mice compromises EC barrier function to maximum at 24 hours and then recovers to baseline over time. 24 hours post LPS administration, freshly made PFK15 (50 mg/Kg) or vehicle control (5% DMSO+45% PEG300+1%Tween 80) were intraperitoneally injected. 30 min before indicated harvest times, mice were retro-orbitally injected with EBA, and lung tissues were harvested for EBA assay. Representative Images of lungs isolated from mice before LPS challenge, 48 hours after LPS treatment, and 48 hours LPS treatment plus 24 hours PFK15 are shown. (**g**) PFK15 decreases the survival rate of LPS-treated mice. C57/bl6 mice were injected with LPS. 24 hours later, they were treated with vehicle control or PFK15, and survival was monitored. The percentage of survival was calculated and plotted. PFK15 alone does not cause mice death, while PFK15 enhances LPS-induced death in mice. Log-ranked analysis shows p<0.001. Source data for (b), (c), (d), (e), (f), and (g) are available in Figure 3 Source Data 1.

**Figure 4.**
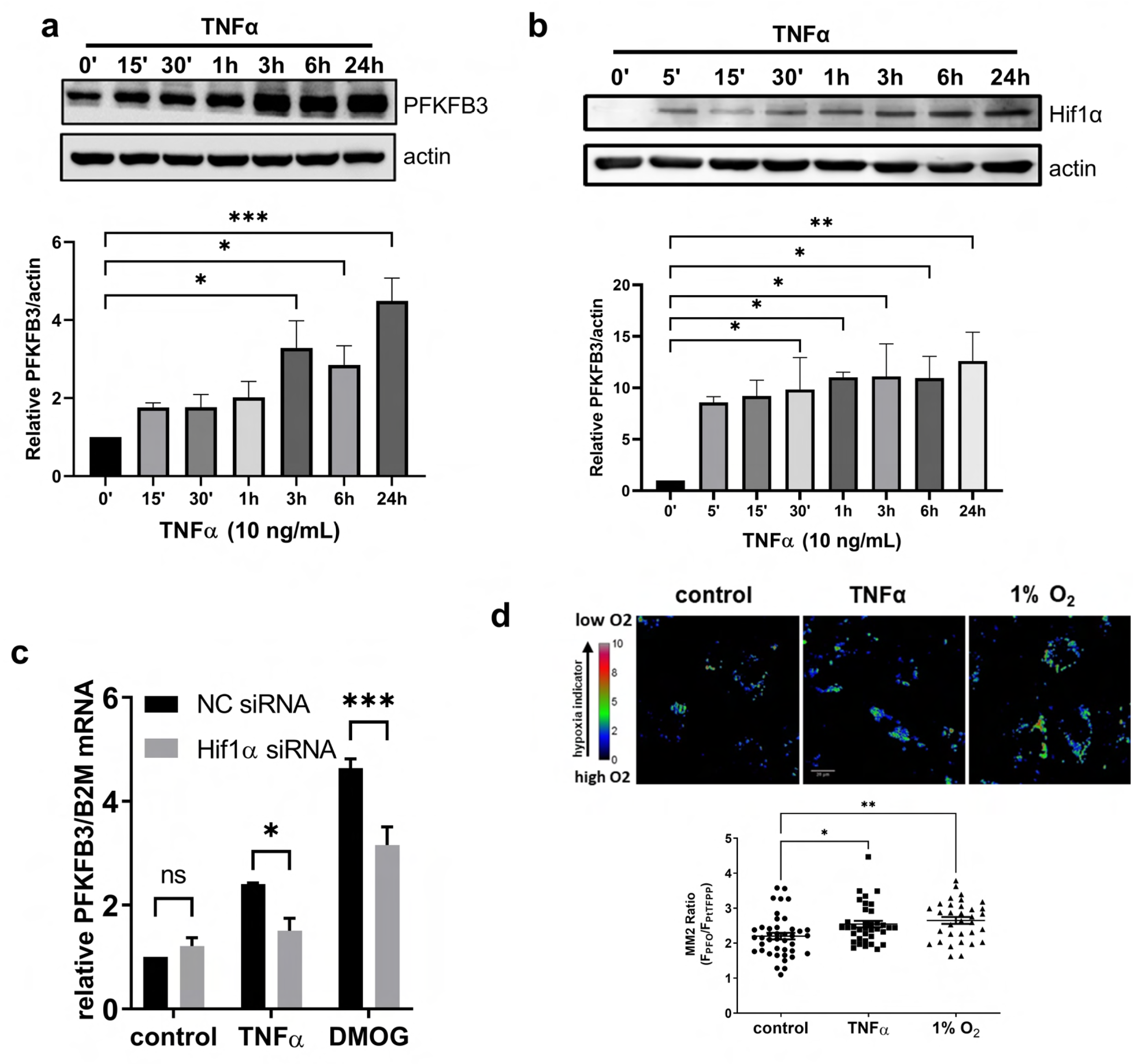
TNFα upregulates PFKFB3 expression via Hif1α-mediated transcription. (**a**) Western blot showing TNFα significantly increases PFKFB3 protein levels in HLMVECs starting at 3 hours after treatment. Both representative image and densitometric quantification performed in ImageJ were shown. (**b**) Western blot showing TNFα significantly increases Hif1α starting 5’ after treatment. Hif1α and actin were quantified using ImageJ. Both representative image and the relative Hif1α/actin are shown. (**c**) Hif1α knockdown decreases TNFα-induced PFKFB3 transcription. HLMVECs were transfected with specific Hif1α siRNAs for knockdown, using non-specific siRNA as a negative control. 48 hours later, transfected cells were treated with TNFα or DMOG (1 mM, to mimic the hypoxia condition) for 3 hours and harvested for RT-PCR. (**d**) Representative MM2 images and quantification showing that TNFα induces intracellular hypoxia. MM2 is a ratio-metric fluorescent hypoxia sensor. MM2-loaded HLMVECs were treated with TNFα or 1% O_2_ for imaging. Raw western blot images for (a) and (b) are available in Figure 4 source data 1 and source data for (a), (b), (c), and (d) are available in Figure 4 source data 2.

PFKFB3 protein levels increase 3 hours following TNFα treatment, providing an explanation for the observed sustained glycolytic activation in ECs, however the upregulation PFKFB3-mediated glycolytic activity occurred within 15 minutes of treatment (**Figure 1**), suggesting that TNFα likely also increases PFKFB3 activity by another complementary pathway. We therefore hypothesized that PFKFB3 was additionally regulated by a more rapid post-translational mechanism, allowing for the rapid upregulation of its activity. TNFα is known to activate the tyrosine kinase Src in tumor cells (Kawai, Tsuji et al. 2002, Rivas, Tkach et al. 2010). Src was shown to activate glycolysis (Zhang, Wang et al. 2017, Pelaz and Tabernero 2022). We hypothesized that Src may mediate TNFα-induced PFKFB3 activation. First, we studied the kinetics of TNFα-induced Src activation by examining the phosphorylation of residue Y416, an indicator of Src activation (Thomas and Brugge 1997). Here we observed that TNFα activates Src as early as 15 min (**Figure 5 Supplementary Figure 1**). We then studied the tyrosine phosphorylation level of PFKFB3 using a phospho-tyrosine specific antibody and found that TNFα exposure rapidly induces PFKFB3 tyrosine phosphorylation within 15 minutes (**Figure 5a**). PP1, a general Src family kinase inhibitor, prevented TNFα-induced PFKFB3 tyrosine phosphorylation (**Figure 5b**). These results suggest a Src family kinase may be responsible for this tyrosine phosphorylation. Moreover, PP1 inhibits TNFα-induced increases in glycolysis (**Figure 5c**), suggesting Src family kinase is involved in TNFα-induced glycolysis activation, possibly via PFKFB3 tyrosine phosphorylation. To verify that Src can directly phosphorylate PFKFB3, we performed an in vitro Src kinase assay. The incubation of human recombinant Src protein with PFKFB3 increased tyrosine phosphorylation of PFKFB3 in a dose-dependent manner (**Figure 5d**), indicating that PFKFB3 is a direct substrate for Src. Moreover, co-expression of constitutively active Src (CA-Src) with PFKFB3 similarly resulted in increased tyrosine phosphorylation. In contrast, overexpression of dominant negative Src (DN-Src) did not increase the phosphorylation level (**Figure 5e**). We also employed an inducible chemo-genetic system to activate Src (Karginov and Hahn 2011). Briefly, iFKBP was inserted in the catalytic domain of Src, generating an inactive form of Src (RapR-Src). When rapamycin is added to the cells, it forms a complex with RapR-Src and co-expressed FRB, leading to a conformational change and subsequent activation of RapR-Src (**Figure 5f**). We performed an immunoprecipitation of RapR-Src and found that PFKFB3 co-precipitated with Src (**Figure 5g**), which results in an increased tyrosine phosphorylation of PFKFB3 (**Figure 5g**). In conclusion, we showed Src activation is sufficient to induce PFKFB3 phosphorylation (**Figure 5g**).

**Figure 5.**
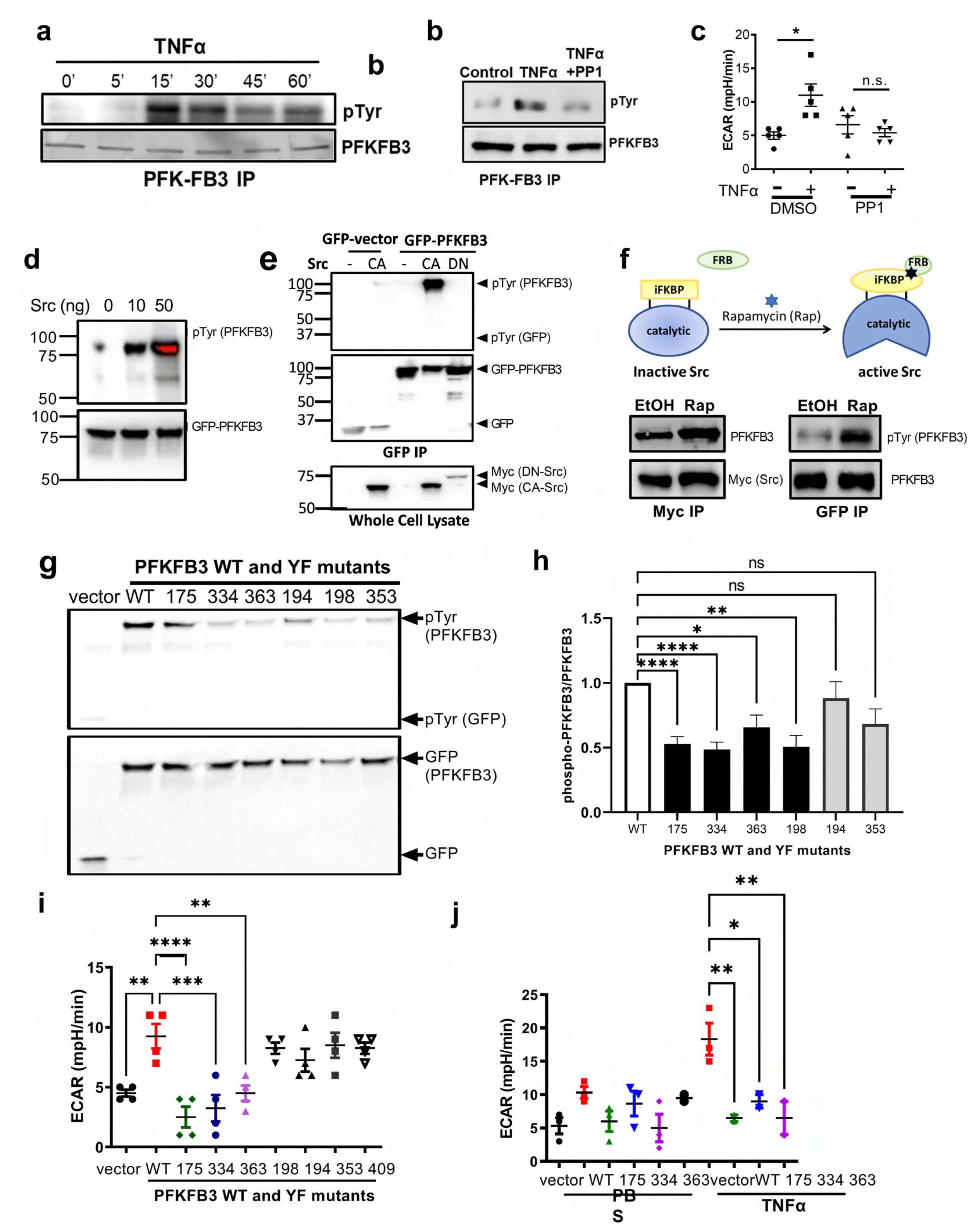
TNFα activates PFKFB3 via Src-mediated tyrosine-phosphorylation. (**a**) Representative western blot images showing that TNFα induces tyrosine phosphorylation of PFKFB3 in a time-dependent manner. Endogenous PFKFB3 was first immunoprecipitated and then subjected to western blot analysis of tyrosine phosphorylation using pan-tyrosine-phospho-specific antibody 4G10. (**b**) Representative western blot showing a selective Src inhibitor PP1 pretreatment inhibits TNFα-mediated PFKFB3 tyrosine phosphorylation. Lysates from treated HLMVECs were immunoprecipitated with PFKFB3, the complex was immuno-blotted with 4G10 and PFKFB3. (**c**) ECAR showing pretreatment with PP1 abolishes TNFα-induced glycolytic activation. HLMVECs were treated with 1 μM PP1 for 30 minutes before TNFα stimulation. ECAR was measured 15 minutes after TNFα treatment. (**d**) In vitro Src kinase assay showed that Src phosphorylates PFKFB3. GFP-PFKFB3 was overexpressed in HEK293T cells and immunoprecipitated using GFP-Trap agarose beads. The PFKFB3 complex was then incubated with the indicated amount of recombinant human Src protein for 10 min. Western blot was used to detect tyrosine-phosphorylation level. (**e**) Representative Western blot images showed that co-expression of constitutively active Src with GFP-PFKFB3 leads to increased tyrosine-phosphorylation in PFKFB3. HEK293T was co-transfected with GFP-PFKFB3 and constitutively active Src or dominant negative Src. GFP-PFKFB3 was pulled down using GFP-trap agarose beads and tyrosine phosphorylation levels were measured by western blot. (**f**) RapR-Src experiments showing that Src binds to and phosphorylates PFKFB3. Schematic of the RapR-Src system shows activation of Src by rapamycin (Rap). An engineered allosteric switch iFKBP domain is inserted to Src kinase domain to generate inactive RapR-Src, Rap triggers the binding of FRB to iFKBP, restoring Src activity. HLMVECs were infected with myc-tagged RapR-Src and FRB. After 1 hour Rap treatment, cell lysate was harvested, Half of the cell lysate were used to immunoprecipitate Src using a myc antibody, and immunoblotted with PFKFB3 antibody to detect its presence in the Src complex, which is shown on the left. The other half cell lysate was used to immunoprecipitate PFKFB3 using PFKFB3 antibody. The WB images are shown on the right. (**g-h**) An in vitro Src kinase assay showed that Src phosphorylates PFKFB3 at Y175, Y334, Y363 and Y198. HEK293T cells were transfected with PFKFB3 WT or its YF mutants, using the GFP vector as a control. PFKFB3 and its mutants were precipitated with GFP-trap agarose beads and used for in vitro Src kinase assay using recombinant Src protein. Representative western blot images of PFKFB3 tyrosine phosphorylation and PFKFB3 after in-vitro Src kinase assay was shown in g. Quantification of PFKFB3 tyrosine phosphorylation over PFKFB3 was shown in h. (**i**) ECAR measurement of PFKFB3 and its tyrosine-phospho-deficient mutants showing that tyrosine phosphorylation at Y175, Y334 and Y363 is important for glycolytic activation. (**j**) ECAR measurement of PFKFB3 and its mutants in HLMVECs in response to TNFα stimulation showing the involvement of tyrosine phosphorylation at Y175, Y334 and Y363 in glycolytic activation by TNFα. HLMVECs were infected with PFKFB3 wild type (WT) and the YF mutants for expression, and they were treated with TNFα and subjected to a seahorse assay for measurement. Raw western blot images are available in Figure 5 Source data 1 and data source for (c), (h), (i), and (j) are available in Figure 5 Source data 2.

NetPhos 2.0 (Blom, Gammeltoft et al. 1999) can predict kinase residues and was used to identify potential Src targets in PFKFB3. 7 residues were identified as potential Src targets (**Figure 5 supplementary Figure 2a**). Using selective mutagenesis, we generated phospho-deficient mutants by mutating each of these seven residues to phenylalanine (YF mutants) and determined how mutations at each site affected total PFKFB3 phosphorylation levels. Using an in-vitro Src kinase assay we found that recombinant human Src protein phosphorylates PFKFB3 at a reduced level when tyrosine at residues 175, 334, 363 and 198 were mutated (**Figure 5h, i**), indicating that these residues are Src targets. To determine whether these residues are phosphorylated in the cellular context, we also induced PFKFB3 phosphorylation by co-expressing PFKFB3 WT/YF mutants and CA-Src in HEK293T cells. Compared to WT, we observed reduced tyrosine phosphorylation in YF mutation at residues 175, 194, 334 and 363 while tyrosine phosphorylation in other YF mutants was not affected (**Figure 5 supplementary Figure 2b, c**). We thus concluded that Src phosphorylates PFKFB3 at residue tyrosine 175, 334 and 336. Additionally, PFKFB3 WT overexpression increased the basal glycolytic activity, which was not observed in YF mutants at 175, 334 and 363 (**Figure 5j**), indicating these tyrosine residues are important for the basal PFKFB3 activity. Moreover, these three mutations also abolished the TNFα induced increase in glycolytic activity (**Figure 5k**). In conclusion, Src-mediated phosphorylation of tyrosine residues 175, 334 and 363 of PFKFB3, is necessary for increasing PFKFB3 activity following TNFα exposure, thus representing a newly identified mechanism of post-translational regulation of PFKFB3 activity.

Using live cell imaging with the ATP sensor PercevalHR (Tantama, Martinez-Francois et al. 2013), we observed local ATP increases in TNFα-stimulated HLMVECs at sites of cell-cell interaction (**Figure 1d**, arrowhead). Additionally, as shown in **Figure 2**, PFKFB3-mediated glycolysis enhances the re-assembly of EC junctions. We thus hypothesized that this effect on re-assembly was due to an augmented regional ATP supply in the subcellular compartments surrounding the AJs at cell-cell contacts. We thus explored how peri-plasma membrane ATP changed in response to TNFα stimulation. The plasma membrane protein stargazin was tagged with iRFP670 and co-expressed with the ATP sensor in HLMVECs. The ATP/ADP ratio was calculated in regions that colocalized with iRFP, indicating the membrane ATP levels (mATP). Consistent with our previous observations, mATP increased in response to TNFα (**Figure 6a, b**). This local ATP increase could have been due to local PFKFB3 activation, PFKFB3 translocation to junction sites, or ATP diffusion. We did not observe obvious PFKFB3 membrane translocation upon TNFα stimulation, excluding this possibility. We next studied whether this spatial regulation of ATP levels is important for restoration of the EC junctions. Although we did not observe increased PFKFB3 at the junctions following TNFα treatment, we hypothesized that overexpressing PFKFB3 specifically at the membrane would increase local ATP, allowing us to evaluate the importance of increased mATP in AJ restoration. We engineered a novel chemo-genetic system to regulate PFKFB3 localization, a Rapamycin-controlled PFKFB3 membrane translocation system (RapT). In this system, PFKFB3 was fused to FKBP12, and FRB was fused to Stargazin. The addition of rapamycin brings PFKFB3 proximal to the plasma membrane by promoting dimerization of the FKBP12 and FRB domains (**Figure 6c**). In HLMVECs expressing the RapT system, upon rapamycin treatment, PFKFB3 was translocated away from the para-nuclear region to colocalize with Stargazin (**Figure 6d**). The addition of Rapamycin significantly increased the Pearson correlation coefficient of PFKFB3 and Stargazin in RapT expressing cells, indicating efficient translocation of PFKFB3 to the membrane (**Figure 6e**). As a parallel approach, we quantified membrane PFKFB3 levels after applying a Stargazin-membrane mask and found increased membrane PFKFB3 levels (Figure **6f**). We then co-expressed PercevalHR and RapT in HLMVECs and studied the effect of controlled membrane translocation of PFKFB3 on mATP levels. Rapamycin induced rapid and sustained PFKFB3 membrane localization starting at 3.5 minutes, followed by an increase in mATP starting at 7 min (**Figure 6g, Figure 6 supplementary data 1**), demonstrating that ATP can be locally modulated by spatial control of PFKFB3. Furthermore, in HLMVECs expressing RapT, rapamycin enhanced the recovery rate of EC barrier function after re-addition of Ca^2+^ (**Figure 6h, Figure 6 supplementary data 2**). The addition of Rapamycin also significantly increased membrane VE-cadherin in cells expressing both RapT constructs, but not in cells over-expressing only the RapT-PFKFB3 component (**Figure 6i, j**). These results indicate that peri-membrane ATP regulates EC barrier function by promoting restoration of VE-cadherin AJ complexes.

**Figure 6.**
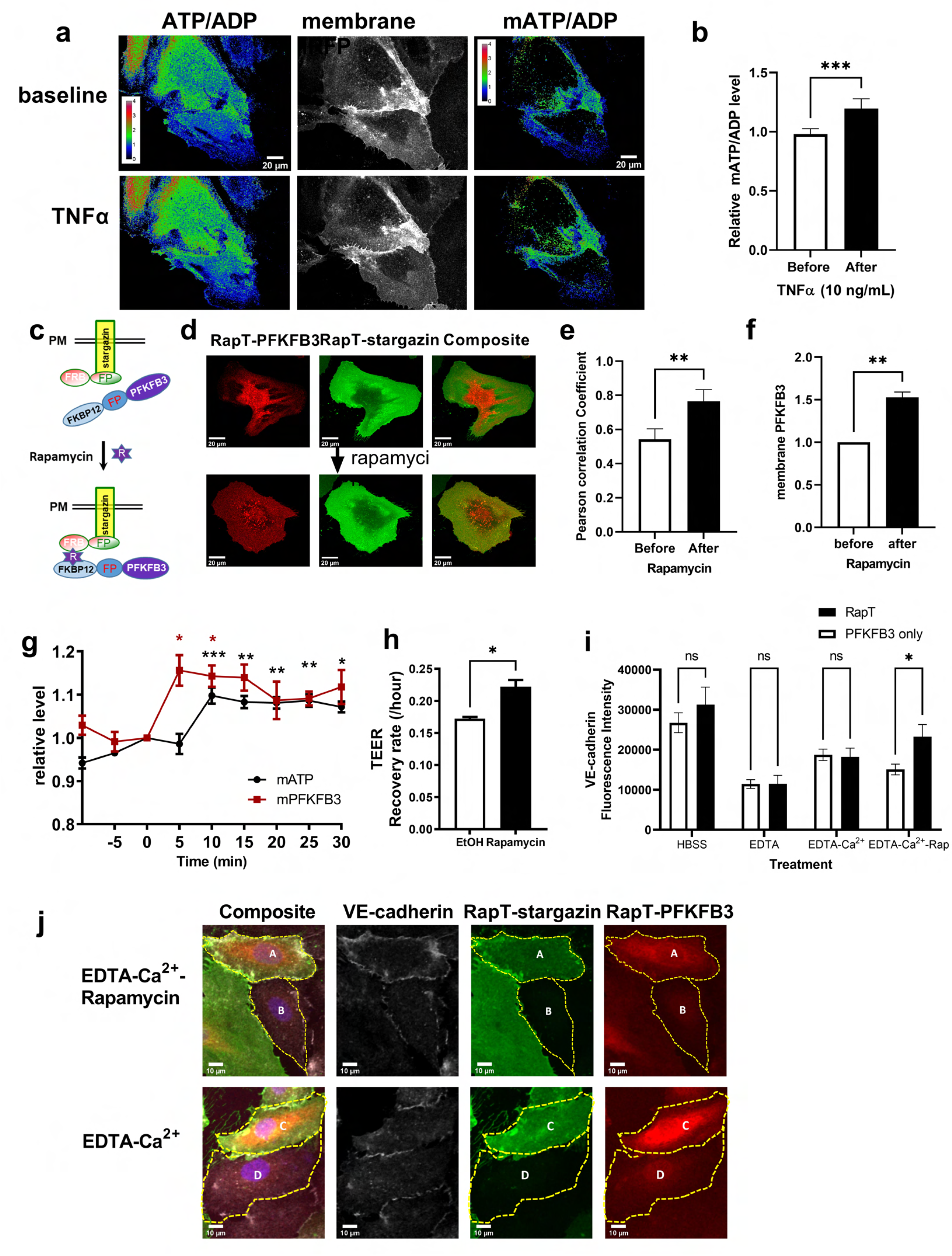
Membrane PFKFB3 enhances mATP production and the subsequent recovery of EC barrier function. (**a**) Representative images showing that TNFα increases membrane ATP/ADP (mATP) levels. ImageJ-Fiji was used to process and analyze images. The first column is the rainbow images of ATP/ADP, the 2^nd^ column is plasma membrane-targeted iRFP670, and the 3^rd^ column is the calculated ATP/ADP that is co-localized with iRFP670, i.e., mATP. The first row is before TNFα treatment, and the bottom row is after TNFα treatment. (**b**) Quantification of TNFα-induced increase of mATP nine cells from 3-independent experiments. (**c**) Schematic diagram of PFKFB3-RapT system: Rapamycin induces PFKFB3 translocation to plasma membrane. Fluorescent protein (iRFP or Venus) and FRB were fused to membrane protein stargazin, while fluorescent protein (mApple) and FKBP12 were fused to nucleus-free (nuc-free) PFKFB3. Rapamycin triggers the dimerization of FKBP12 and FRB, bringing PFKFB3 from cytosol to the plasma membrane. (**d**) Representative images showing that rapamycin induces PFKFB3 translocation to plasma membrane. The top three images are before treatment: stargazin-Venus-FRB, nuc-free PFKFB3-mApple-FKBP12, and composite. The bottom images are the corresponding images after treatment (Rapamycin 1μM, 5 minutes). (**e**) Rapamycin increases the Pearson correlation coefficient of PFKFB3 with stargazin. The ImageJ plugin JACoP was used to calculate the Pearson correlation coefficient. (**f**) Rapamycin increases membrane PFKFB3. Stargazin-iRFP images were converted to a binary image (“1” for iRFP expression and “0” for no iRFP expression), the membrane PFKFB3 is calculated by multiplying PFKFB3 with the binary iRFP images. (**g**) Quantification to show the kinetics of Rapamycin-induced mPFKFB3 and mATP increase. mATP and mPFKFB3 were calculated, and then normalized to baseline and plotted. The results shown are the average of eight cells from three independent experiments. (**h**) Quantification of resistance recovery rate of TEER showing Rapamycin-induced PFKFB3 plasma membrane translocation enhanced recovery of EC barrier function (n=3). HLMVECs in TEER plates were first treated with 1mM EDTA to disrupt junction, then rapamycin (0.5 μM) or Ethanol (0.05%, vehicle control) was added simultaneously with Ca^2+^ to study the effect of membrane PFKFB3 on the recovery of EC barrier function. The recovery rate was calculated and plotted. (**i**) Quantification of membrane VE-cadherin fluorescence in RapT (cells expressing both RapT constructs), PFKFB3-only (Single Positive cells expressing RapT-PFKFB3, but not RapT-stargazin) showing that rapamycin enhances membrane VE-cadherin localization in RapT cells but not in PFKFB3-only cells. HLMVECs were infected with lentivirus expressing stargazin-Venus-FRB and PFKFB3-mApple-FKBP12. seven days later, cells were treated with EDTA for 15 minutes, followed by adding Rapamycin (1μM) and/or Ca^2+^ (1mM) for 45 minutes. Cells were then fixed and stained for VE-cadherin and DAPI to stain the nucleus (blue). (**j**) Representative images of HLMVECs treated with either EDTA+Ca^2+^ (bottom row) or EDTA+Ca^2+^-Rapamycin (top row). Cells A and C expressed both RapT constructs while cells B and D only expressed RapT-PFKFB3. The composite images, VE-cadherin (white), RapT-stargazin (green) and RapT-PFKFB3 (red), RapT-stargazin (green). Data source for (b), (e), (f), (g), (h) and (i) are available in Figure 6 Source data 1.

## Discussion

In this study, we have uncovered the importance of PFKFB3-mediated glycolytic ATP availability for the restoration of endothelial adherens junctions. The conventional view of inflammatory signaling is that it leads to the disruption of AJs and thus promotes vascular leak and edema formation. However, we show that inflammatory signaling also concomitantly generates a metabolic milieu for increased glycolytic ATP generation which is essential for the restoration of the endothelial barrier, thus serving as an important adaptive mechanism for the resolution of the initial injury (**Figure 7**). It suggests a finely tuned coordination of barrier disruption and barrier restoration with a key role for the Src-PFKFB3 axis.

**Figure 7.**
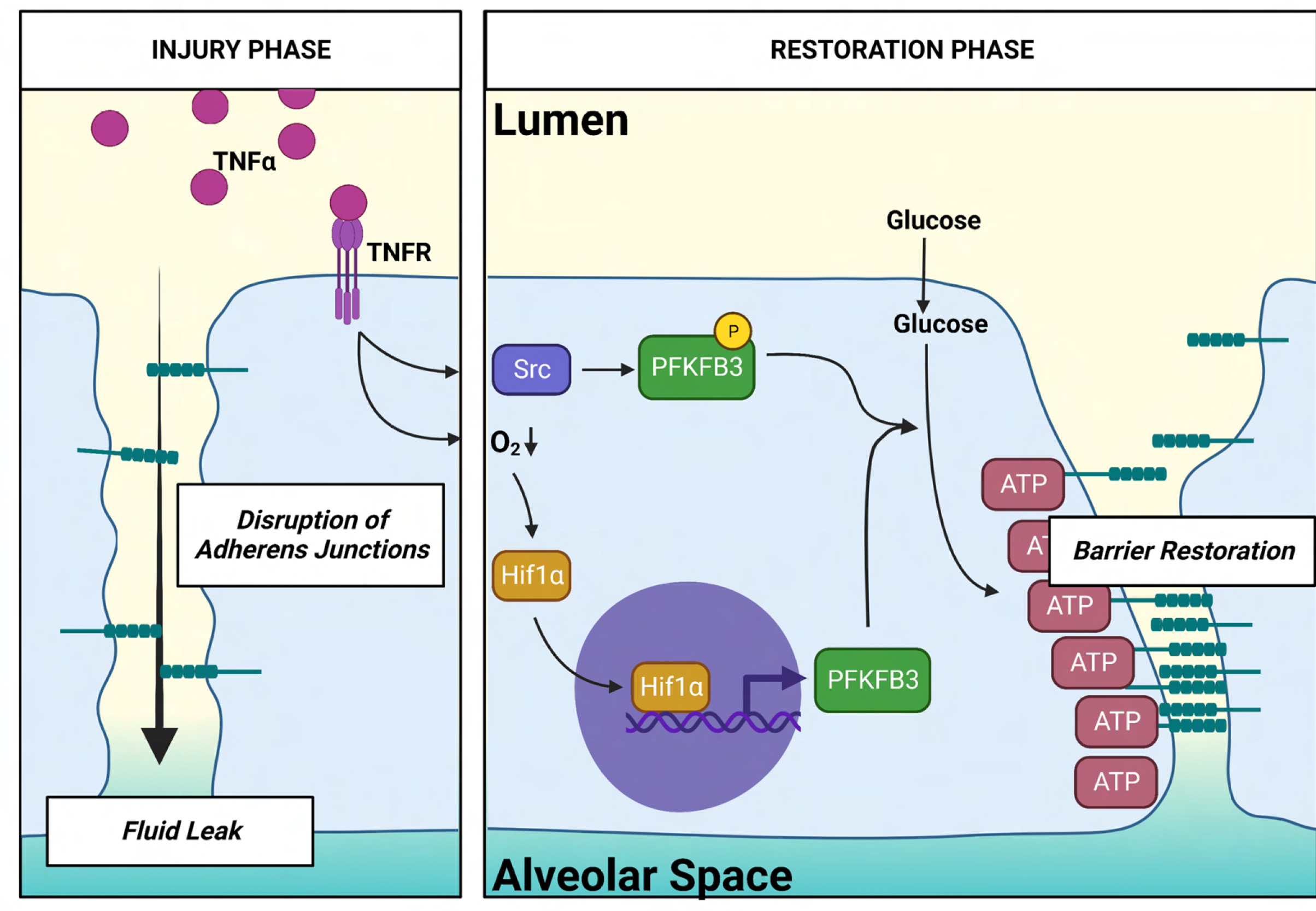
Transcriptional and post-translation regulation of PFKFB3 increases intracellular ATP, which accelerates the AJs re-assembly during resolution. During the injury phase, inflammatory stimuli induce disruption of adherens junctions, thus resulting in fluid leak. In anticipation of the subsequent restoration, the same inflammatory stimuli concomitantly activate PFKFB3 via Src and Hif1α, resulting in ATP increase near AJs and promoting EC barrier restoration by accelerated AJs re-assembly.

Under homeostatic conditions, the PFKFB3 inhibitor PFK15 dose-dependently disrupted EC barrier function and junction tension. These results suggest that glycolytic ATP is needed to maintain VE-cadherin at the AJs. As VE-cadherin is actively trafficked between the plasma membrane and intracellular compartment (Giannotta, Trani et al. 2013), it is possible that glycolytic ATP is required for dynamic VE-cadherin trafficking. In tumor ECs, the glycolysis inhibitor 3PO has biphasic dose-dependent effects with a low dose of 3PO leading to tumor vessel normalization (Cantelmo, Conradi et al. 2016), and higher doses causing vessel disintegration (Conradi, Brajic et al. 2017). These results indicate the possibility that glycolytic ATP may be involved in both the internalization and the recycling of cadherins to AJs. Here, we observed that restoration of AJs is delayed when PFKFB3 is inhibited while it is enhanced when PFKFB3 is overexpressed. Our results complement the observations that another glycolytic enzyme, PKM2, is also required for pulmonary microvascular barrier function (Gomez-Escudero, Clemente et al. 2019). Importantly, we showed that following inflammatory injury, PFKFB3 promotes restoration of EC barrier function by accelerating re-assembly of VE-cadherin, which is consistent with reports that CD31-induced glycolytic activation enhances endothelial barrier recovery (Cheung, Fanti et al. 2020).

We observed that PFKFB3 was activated downstream by TNFα signaling in a two-pronged manner, rapidly through post-translational modification by Src as well as in a more sustained manner via transcriptional upregulation of PFKFB3 by Hif1α. The oncogenic kinase Src has been previously shown to increase glycolysis (Rempel, Mathupala et al. 1996). Here we have uncovered tyrosine phosphorylation of PFKFB3 by Src as a novel mechanism by which Src regulates glycolysis. We identified 4 tyrosine residues on PFKFB3 (residues 175, 194, 334 and 363) which are targets of Src activity. One of the residues identified as being phosphorylated here, Y194, was recently shown to be phosphorylated by c-Src to enhance glycolysis and promote tumorigenesis (Ma, Zhang et al. 2020). However, we did not observe that this residue regulates glycolysis in ECs during inflammation. Instead, we found that phosphorylation of the other 3 residues plays a role in regulating endothelial glycolysis. Of note, both Y363 and Y334 are in the phosphatase domain of PFKFB3. Notedly, Y363 has been shown to be a critical residue mediating the binding of the 6-phosphate moiety of either F-2,6-biphosphate or F-6-phosphate (Kim, Manes et al. 2006). Thus, phosphorylation of these two residues may interfere with the phosphatase activity, thus resulting in increased F-2,6-biphosphate levels and thereby activating glycolysis.

We also explored the spatial regulation of ATP by PFKFB3 using a novel engineered rapamycin-membrane translocation system (RapT-PFKFB3), as TNFα-stimulated HLMVECs specifically increased ATP in the proximity of the plasma membrane. Although we were not able to detect endogenous PFKFB3 translocation to the cell-cell junctions in response to inflammatory activation, membrane recruitment of PFKFB3 in our RapT system increased local ATP and enhanced the recovery of EC barrier function. These results suggest that availability of ATP at the junctions is a determining factor in the rate of EC barrier restoration. Further study is needed to determine the mechanism of this localized ATP production, which could be mediated by regional post-translational activation of PFKFB3, rather than through translocation of the enzyme. Additionally, the recent development of engineered optogenetic (LightR) allosteric switches (Shaaya, Fauser et al. 2020) could allow for precise spatiotemporal control of enzyme activity *in vivo* and *in vitro*. Additionally, our use of fibronectin-coated PDMS stamps coupled to a VE-cadherin tension sensor allowed for the measurement of tension across individual junctions and thus avoided confounding tension forces that are present in monolayers where cells are in contact with several neighbors. Our approach could be expanded to study more global effects of PFKFB3 inhibition on cell processes such as traction and cytoskeletal orientation relative to isolated junctions.

Recent studies are identifying the critical role of compartmentalized metabolic shifts in the regulation of cellular junction homeostasis and integrity such as the observation that the glucose transporter Glut1 is recruited to junction sites to meet the energy requirements of cytoskeleton remodeling responses to shear stress (Salvi, Bays et al. 2021). Localized expression of the glucose transporter in the front end of invasive cells provides ATP burst to breach basement membrane barrier (Garde, Kenny et al. 2022). In this study we show that PFKFB3, a glycolysis regulatory enzyme that is downstream of the Glut1 transporter, is required for reassembly of adherens junction and that post-translational modification of this essential metabolic enzyme is a means by which restoration of cellular barrier integrity can be controlled. Enhanced regional ATP production at the junctions accelerates adherens junction re-assembly and barrier restoration, thus underscoring the importance of localized ATP availability. Engineered constructs that allow for the spatial control of localized ATP generation such as the RapT-PFKFB3 combined with localized compartment-specific assessment of ATP levels are essential for understanding the roles of localized ATP generation and these insights may lead to novel therapeutic approaches that focus on localized metabolic shifts to accelerate repair of barriers that are disrupted during disease processes.

## Materials and Methods

### Materials

Chemicals were obtained from Sigma-Aldrich, and cell culture reagents were obtained from Fisher Scientific unless otherwise specified. PFK15 was obtained from Cayman Chemicals, and human recombinant TNFα from R&D systems. Small interfering RNAs (siRNAs) targeting human PFKFB3, human Hif1α and non-specific control siRNA, siRNA transfection reagent, and polybrene were obtained from Santa Cruz Biotechnology.

### Plasmid constructs

Human PFKFB3 tagged with N-terminal GFP was cloned into the viral plasmid pWPXL (Addgene #12257). A PFKFB3 mutant (nuc-free PFKFB3) losing nuclear localization was generated by mutating the lysine residues at 472 and 473 to alanine residues (KK472/473AA) via the QuickChange site-directed mutagenesis kit (Stratagene). The same approach was used to generate tyrosine phospho-defective PFKFB3 at the indicated residues by replacing tyrosine with phenylalanine. To mark plasma membrane, the membrane protein Stargazin was fused to a fluorescent protein iRFP670 at its C-terminus, and then cloned to a viral plasmid Lego-iV2 (Addgene #27344). To generate the RapT system, nuc-free PFKFB3 was used to ensure free accessibility for plasma membrane translocation upon rapamycin stimulation. mApple was first fused to nuc-free PFKFB3 at its C-terminus, and then fused to FKBP12 at its N-terminus. This whole fragment was then cloned into the viral plasmid pLJM1-EGFP (Addgene #19319). Stargazin was fused to Venus-FRB (Q157cpFRB), the whole segment was cloned into pWPXL. To measure the ratio of ATP to ADP concurrently with Rapamycin-mediated PFKFB3 membrane translocation in the RapT system, another membrane-localized FRB (stargazin-iRFP670-FRB) was also generated by inserting FRB into Lego-iV2-stargazin-iRFP670 via the Q5 site-directed mutagenesis kit (NEB). The FRET-based VE-cad-TS (Addgene Plasmid# 45848) was used to measure tension on VE-cadherin bonds. For expression in endothelial cells, the construct was cloned into the PLJM1-EGFP lentiviral vector (Addgene, Plasmid #19319). The inserts were amplified by PCR, using the Q5 Polymerase (New England Biolabs) and ligated into the digested PLJM1 vector at AgeI and EcoRI restriction sites. All cloning changes listed above were confirmed by Sanger sequencing. Further confirmation also included the correct localization as visualized under fluorescent microscope and successful expression by western blot.

### Cell culture

Human lung microvascular endothelial cells (HLMVECs, Cell Application Inc, cat# 540-05a) were cultured in flasks coated with 2% gelatin using endothelial basal medium 2 (EBM2, Lonza) supplemented with bullet kit additives plus 10% fetal bovine serum. HLMVECs between passages 3 and 10 were used for experiments. HEK293T (ATCC, Cat# CRL-3216) was cultured in DMEM supplemented with 10% FBS and 1% Pen-Strep. 293AD cells were cultured in DMEM supplemented with 10% FBS, 0.1mM NEAA, 2mM L-glutamine and 1% Pen-strep.

### siRNA transfections

HLMVECs were transfected with siRNAs (si-PFKFB3, si-Hif1α, and nonspecific scramble control siRNA) using siRNA transfection reagent (Santa Cruz Biotech, Cat# sc-29528) according to the manufacturer’s instructions. 48 hours later, cell lysates were harvested to confirm the successful knockdown of the protein of interest. HLMVECs at 48 hours post-transfection were used to perform subsequent ECAR, Western or RT-PCR experiments.

### Virus generation

To generate lenti-virus including PFKFB3 and its mutants, PFKFB3 KKAA-mApple-FKBP12, Stargazin-Venus-FRB, stargazing-iRFP-FRB, ATP sensor FUGW-perceval HR, and VE-Cadherin tension sensor, HEK293T cells were co-transfected with VSV-G (the envelop expressing plasmid, addgene, #12259) and psPAX2 (the virus packaging plasmid, addgene, #12260) using Jetprime (Polyplus, Cat# 101000046) based on the manufacturer’s protocol. 48 hours and 72 hours post transfection, supernatants were harvested. Viral particles were precipitated using Lenti-X maxi-purification kit (Takara, Cat# 631233) based on manufacturer’s protocol. HLMVECs were transduced with lentivirus at a multiplicity of infection of 5-20, together with polybrene (final conc. 4 μg/mL). Expression was generally observed 1-7 days after infection.

To amplify adenovirus including RapR-Src and Myc-FRB (gift from Dr. Karginov at UIC), 293AD (gift from Dr. Karginov at UIC) cells were first infected with adenovirus. 48 hours after infection, cells were harvested, and the viruses were purified using Adeno-X maxi purification kits (Takara, Cat# 631533) based on the manufacturer’s protocol. HLMVECs were infected with RapR-Src and Myc-FRB the day before experiments. 16 hours later, cells were ready for experiments.

### RT-PCR

Treated cells were harvested at the indicated time for mRNA isolation. mRNA was extracted using the TRIzol reagent following manufacturer’s TRIzol RNA isolation protocol. 1-2 μg RNA was then converted to cDNA using high-capacity cDNA reverse transcription kits (Applied Biosystems). cDNA (1 ng/μL) was then subjected to quantitative PCR using FastStart Universal SYBR Green Master reagents (Rox) ABI7500 Real Time PCR system (applied biosystems). Relative gene expression was calculated using the 2^-ΔΔt^ method against the internal control B2M.The sequence of the primers we used are: human PFKFB3 forward (5’ ATTGCGGTTTTCGATGCCAC 3’), human PFKFB3 reverse (5’ GAACGTCGAAAAGAAAAGTCTCG 3’), human Hif1α forward (5’ GAACGTCGAAAAGAAAAGTCTCG 3’), human Hif1α reverse (5’ CCTTATCAAGATGCGAACTCACA 3’), human B2M forward (5’ GGTTTCATCCATCCGACATT 3’), human B2M reverse (5’ ATCTTTTTCAGTGGGGGTGA 3’)

### Co-immunoprecipitation (Co-IP) and Western blot

For Co-IP studies, treated cells were lysed in cell lysis buffer (10mM Tris HCl, 150mM NaCl, 1mM EDTA, 1mM EGTA, 1% Triton X-100, 0.5% NP40 supplemented with protease and phosphatase inhibitor cocktail). Specific protein complex was immunoprecipitated by incubating the lysates with agarose A/G beads and a specific antibody at 4°C overnight. After washing with cell lysate buffer with 1mM PMSF, beads with bound complex in 2X loading buffer were applied to SDS-PAGE for western blot. One tenth of the total cell lysates were used as input controls.

For Western blot analysis, treated cells were lysed in RIPA buffer supplemented with protease- and phosphatase-inhibitor cocktail. Lysates with equal amounts of total protein were loaded onto 8%-12% polyacrylamide gels, separated by electrophoresis, and transferred to nitrocellulose membranes. After blocking in 5% BSA at RT for 1 hour, the membranes were probed with primary antibody overnight at 4°C and secondary antibody 1 hour at RT before developing. The primary antibodies were: PFKFB3 (Proteintech, Cat#13763-1-AP), Hif1α (Cayman chemicals, Cat#10006421), 4G10 (Millipore, Cat# 05-321), Src antibody (Santa Cruz Biotechnology, Cat# sc-18), phosphor-Src (Y416, Cell signal, Cat# D49G4), Myc (Santa Cruz Biotechnology, Cat# sc-40), GFP (Clontech), and β actin (Sigma, Cat# A5316). The intensity of protein level was quantified using Image J software.

### In vitro Src kinase Assay

HEK293T cells were transfected with GFP-PFKFB3 and its YF mutants for overexpression. 24 hours later, cell lysate was lysed in RIPA buffer (10 mM Tris/Cl pH 7.5, 150 mM NaCl, 0.5 mM EDTA, 0.1% SDS, 1% Triton X-100, 1% deoxycholate) and the lysates were used for PFKFB3 immunoprecipitation using GFP-TRAP agarose beads (Proteintech, cat# gta) according to the manufacturer protocol. The immunoprecipitants were washed with kinase buffer (25 mM HEPES pH7.5, 5 mM MgCl_2_, 0.5mM EDTA, 0.005% BRIJ-35) for two times and used to performed in vitro Src kinase assay. The PFKFB3 immunoprecipitant in 20 μL Kinase buffer and 200 uM ATP was mixed with 20uL of human recombinant protein (20 ng, active motif, cat# 81115) and incubated at 37°C for 10 min before stopping the reaction by the addition of 60uL of 2X Laemmli protein sample buffer. The reaction was loaded to SDS-PAGE and immune-blotted with 4G10 and GFP to access the tyrosine phosphorylation level in GFP-PFKFB3.

### Trans-endothelial electrical resistance (TEER) measurement

TEER is a non-invasive method that electrically monitors cell permeability. Thus, ECIS was used for *in vitro* study of barrier function of endothelial cells (ECs) by measuring their resistance. In brief, HLMVECs were plated to gelatin-coated electrode arrays (8W10E+ PET, ECIS) at a density of 10^5^ cells/ml. On the day of experiments, cells were first checked under the microscope to ensure that the cells were at close to 100% confluency. TEER was carried out in HBSS at 37°C and 5% CO_2_. After an initial 1.5 hours equilibrium incubation, changes in electrical resistance were recorded over the entire experiments. With stable baseline recorded for a minimum of 15 minutes, cells were treated with the indicated reagents. PFK15(1-10 μM) was used to inhibit PFKFB3 in the study of PFKFB3 in both maintaining and recovering barrier function. 1mM EDTA was used to chelate extracellular Ca^2+^ and disrupt EC junctions. And 1mM Ca^2+^ was added to restore the EC junctions. TEER was normalized to the baseline to account for variations from well to well and from batch to batch.

### Preparation of polydimethylsiloxane (PDMS) stamps with micro contact printed fibronectin patterns

PDMS stamps were created to microcontact print patterns of tangential 1600 μm^2^ circles of fibronectin on 40 kPa gels. To create the PDMS stamps, 10:1 Sylgard 184 Silicone Elastomer Base, part B (Sylgard^tn^ 4019862) and Sylgard 184 Silicone Elastomer Curing Agent, part A, respectively. The mixture was poured over a negative master and put into a decanter for 1 hour, to remove bubbles. Then the mixture was cured for 15 minutes in a 108° C oven. The PDMS was peeled from the master and cut into individual stamps (Qin, Xia et al. 2010). PDMS stamps were submerged in ethanol and sonicated for 5 minutes, and plasma treated (Harrick Plasma PDC-32G) to make the stamps hydrophilic. Fibronectin (0.1 mg/mL) was pipetted onto the stamps and incubated for 1 hour, to allow fibronectin to adsorb on the stamps. Fibronectin was immobilized on Sulfo-SANPAH-activated (25 mg/μl), 40 kPa gels by aspirating the PBS from the stamp and then stamping the adsorbed protein onto the activated gels. Gels were incubated at 37° C before the stamps were removed. The gels with printed fibronectin patterns were then submerged in PBS and stored overnight at 4° C.

### Immunofluorescence and confocal microscopy

HLMVECs cultured on coverslips were fixed with 4% paraformaldehyde, permeabilized with 0.1% Triton X-100 and blocked with 5% goat serum for 1 hour at room temperature. Slides were probed with primary antibodies overnight at 4°C and fluorescence-conjugated secondary antibodies for 2h at RT. Images were taken with a Zeiss Laser Scanning microscope 510 META (LSM510 META).

Live-cell imaging was performed using a Zeiss LSM710 BIG microscope equipped with a Plan-Apochromat 63x/1.40 Oil DIC objective (Zeiss), and GaAsP and PMT detectors, at 37°C, 5% CO_2_. The day before imaging, cells were plated on gelatin-coated glass-bottom dishes. Multiple positions were imaged using the autofocus function. Before treatments, cells at multiple positions were imaged for 10 minutes to establish the baseline. For intracellular ATP measurement, PercevalHR was excited with 488/10 nm (F488, responding to ATP binding) and 420/30 nm (F420, responding to ADP binding), emission was collected at 500/50. For measuring PFKFB3 translocation by rapamycin treatment, mApple (Ex 568, Em 592) was excited at 561 nm and emission was measured at 575/610, iRFP670 (Ex 643, Em 670) was excited at 633 nm and emission was measured at 637/735, Venus (Ex 515, Em 528) was excited at 514 nm and emission was measured at 518/89.

For measuring tension across VE Cadherin junctions, HLMVECs expressing the VE-cad-TS were imaged at 40x with a Zeiss Axiovert 200M wide field microscope. Regions of interest were imaged at each junction with a CFP (filter set 47, #000000-1196-682, excitation: BP 436/20, emission: 480/40) filter and a FRET (CYP/YFP filter set 48, 000000-1196-684, excitation: BP 436/20, emission: 535/30) base pair filter. Intensities determined with the CFP filter, FRET filter, and background were quantified and analyzed in ImageJ. The background of each was subtracted for both filters, using ImageJ and measuring the background-subtracted intensity at the junctions. FRET/CFP ratios were calculated using ImageJ. As an increase in tension results in a decrease in FRET, the inverse ratio, CFP/FRET was graphed, such that changes in CFP/FRET correspond with the change in tension.

ImageJ/Fiji software package was used to analyze the images. For analysis of the ATP sensor, images from the channel F488 were first subtracted with background. Then fluorescence ratio images were created by pixel-by-pixel division between channel F488 and F420 (F488/F420). Pseudo-colored images were generated to present the actual fluorescence ratio. To calculate membrane ATP/ADP, the plasma membrane iRFP670 images were first converted to binary images (lower threshold is 0 while higher threshold is 1). These images were then multiplied pixel-by-pixel with the ATP/ADP fluorescence ratio to calculate membrane ATP/ADP. A similar approach was used to calculate mPFKFB3.

### Glucose metabolism measurement using Seahorse

Extracellular Acidification Rate (ECAR) measured by seahorse is an indicator of glycolysis. HLMVECs were infected with PFKFB3 WT or its mutants for overexpression and used for experiments 48 hours post-transfection. HLMVECs were transfected with PFKFB3 siRNA and used for experiments 72 hours post-transfection. These HLMVECs were seeded in an XF24 cell culture plate at 100,000 cells/well and were treated as indicated. At the indicated time, treated cells were then equilibrated in seahorse XF base medium supplemented with 1g/L glucose at 37°C in non-CO_2_ incubator for 1 hour prior to analysis using a Seahorse XF24 Extracellular Flux Analyzer (Seahorse Bioscience).

Glycolysis stress kit was used to study acute ECAR change upon TNFα stimulation using a seahorse eXF96 extracellular flux analyzer (Seahorse Bioscience). HLMVECs were seeded in an XF96 cell plates at 25,000 cells/well for analysis the following day. After equilibration in seahorse XF base medium for 1 hour. Chemicals were added to probe glycolysis activity. After injection, three measurements were performed unless specified. Firstly, ECAR of cells in XF base medium was measured for the non-glycolytic activity. Secondly, ECAR of cells after injection with glucose (10 mM) were measured for glycolysis activity under basal conditions. Thirdly, they were stimulated with TNFα (10ng/mL) and ECAR was measured for 150 min (25 measurements). Fourthly, ECAR of cells after injection with oligomycin (1 μM) revealed cellular maximum glycolytic capacity. Oligomycin inhibits ATP synthase and shifts the energy production to glycolysis. Lastly, decreased ECAR of cells after injection with 2-deoxy-glucose (2-DG) confirmed previous ECAR change is due to glycolysis. 2-DG inhibits glucose hexokinase and subsequent glycolysis. To study the effect of PFKFB3 inhibition by PFK15 on TNFα-induced ECAR increase, PFK15 (10 μM) was added at the same time of TNFα treatment. At the same time, Oxygen consumption rate (OCR) was collected to explore the effect of TNFα treatment on mitochondrial respiration activity. Both ECAR and OCR were analyzed using WAVE software provided by Seahorse Bioscience.

### Animal procedures

Animal procedures were approved by and performed in accordance with guidelines by UIC animal care and use committee. For retro-orbital injection, mice were anesthetized with 2% isoflurane inhalation at flow rate of 0.6 liter/min. For endpoint experiments, mice were anesthetized with intraperitoneal administration of a mixture of Ketamine (100mg/Kg), Xylazine 434 (2mg/Kg) and acepromazine (2 mg/Kg) in saline solution.

### Mice treatment

C57BL/6J male mice (8 weeks) were purchased from Jackson Laboratories and were accommodated for two weeks before performing any experiments. PFK15 (5 mg/mL) was prepared by adding solvents to reach the final concentration of 5% DMSO, 45% PEG300, 1% Tween80 in water solution. C57bl/6 mice were first challenged with LPS (15-20 mg/Kg as indicated) or saline control via IP injection. 24 hours post LPS challenge, the mice were randomly separated in two groups, and they were intraperitoneally injected with either PFK15 (50 mg/Kg) or vehicle control (5% DMSO, 45% PEG300, 1% Tween80).

### Evans blue and Album (EBA) assay was used to assess the lung vascular permeability

The lung vascular permeability was measured based on the accumulation of blood BSA-conjugated Evans blue in**to** the lung as described previously (Shigematsu, Koga et al. 2016). 40 minutes prior to harvesting the lung, treated mice were injected with EBA (4% Albumin, 0.5% Evans blue in PBS) to a final concentration of EBA at 20mg/kg via retro-orbital injection. The lung was perfused with PBS to clear EBA in the blood. Next, the lung was removed and grinded in 1ml PBS. Evans blue was extracted by adding 2 ml formamide and incubating the mixture for 18 hours at 55–60°C. Supernatant was separated by centrifugation. The optical density of the supernatant was determined spectrophotometrically at 620 nm (Evans blue) and 740 nm (hemoglobin correction). Corrected A620 was calculated based on the formula: A_620_ (corrected)=A_620_-1.426*A_740_+0.03. The amount of EBA leaked into the lung was calculated using the corrected A_620_ and the standard curve. Body weight was shown to be proportional to the lung size (Parker and Townsley 2004) and was then used to normalize EBA extracted from the lung

### Data collection and Analysis

All experiments are performed using three or more biological replicates. Data was represented as mean ± SEM. ANOVA analysis or t-test were used for the statistical analysis, depending on the sample numbers. p<0.05 (*), p<0.01 (**), p<0.001 (***).

## Acknowledgments

We thank Dr. Ankit Jambusaria for analyzing the promoter region of human PFKFB3, Jake Matsche for technical input on in vitro Src kinase assay, and Dr. Glenn Marsboom for comments and feedbacks on the paper. This project was supported by NIH grants P01HL060678, P01HL160469, R01GM127554, R21CA223915, T32HL007829, T32HL139439 and AHA grant 18PRE34070092.

## Figure Supplement Legends

**Figure 1 Supplementary Figure 1.**
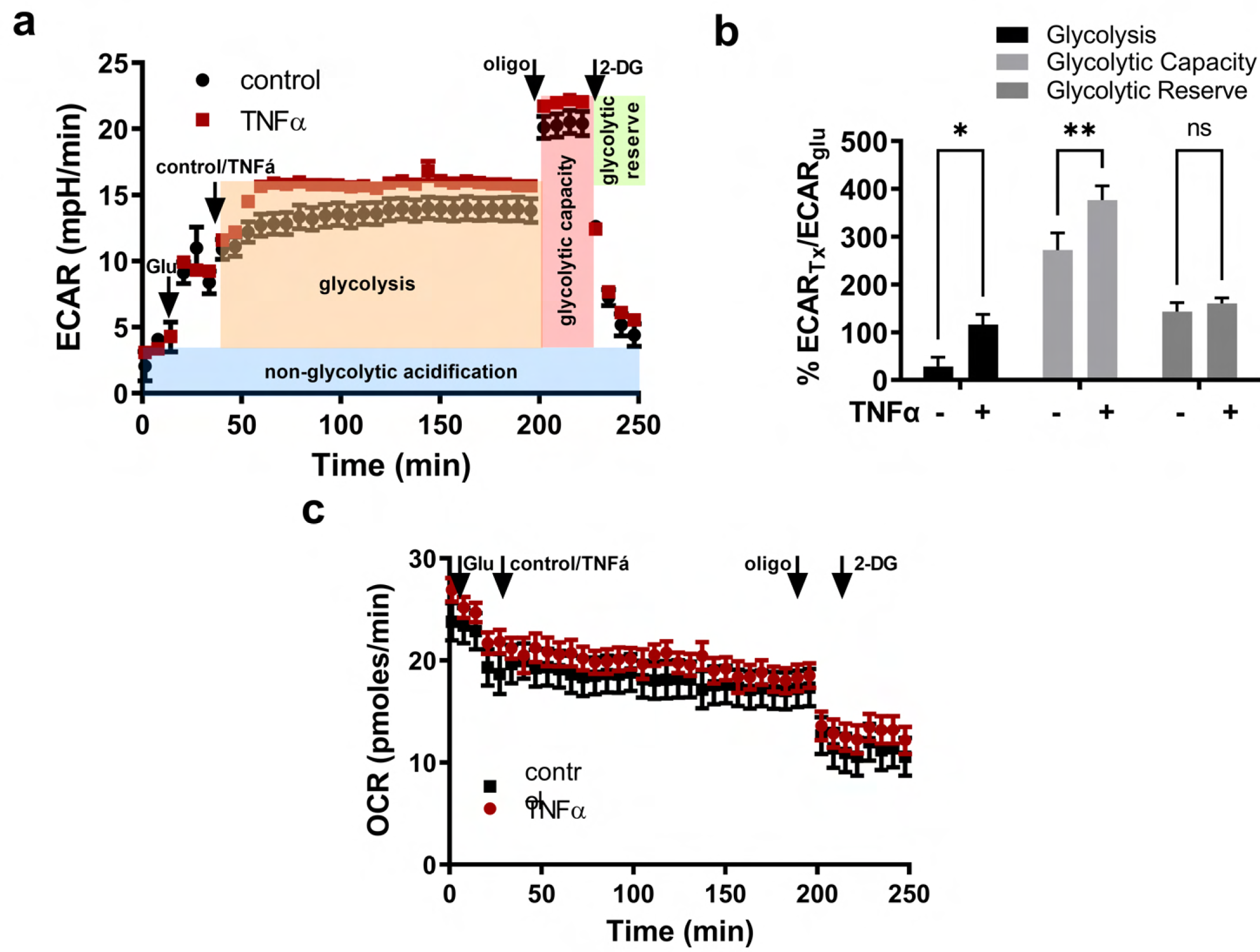
TNFα activates glycolysis but not mitochondrial respiration. (**a**) A representative kinetic ECAR measurement of HLMVECs to glucose (1.0 g/L), TNFα (red square)/ its vehicle control (black triangle), oligomycin (1 μM) and 2-DG (100 mM). TNFα induces a rapid and sustained increase in ECAR in HLMVECs. As a control, glucose increases ECAR and 2-DG decreases ECAR. (**b**) Detailed quantified analysis of the effect of TNFα on glycolytic parameters. Glycolysis, glycolytic capacity, and glycolytic reserve are calculated as the difference between ECAR measurement 11 and 4, 32 and 4, 32 and 11 respectively, and normalized by the glucose-induced glycolysis, which is the difference of measurement 4 and 3. (**c**) A representative kinetics OCR measurement showing that TNFα does not increase mitochondrial respiration. Source data for (a) (b), and (c) are available in Figure 1 Source Data 2.

**Figure 1 Supplementary Figure 2.**
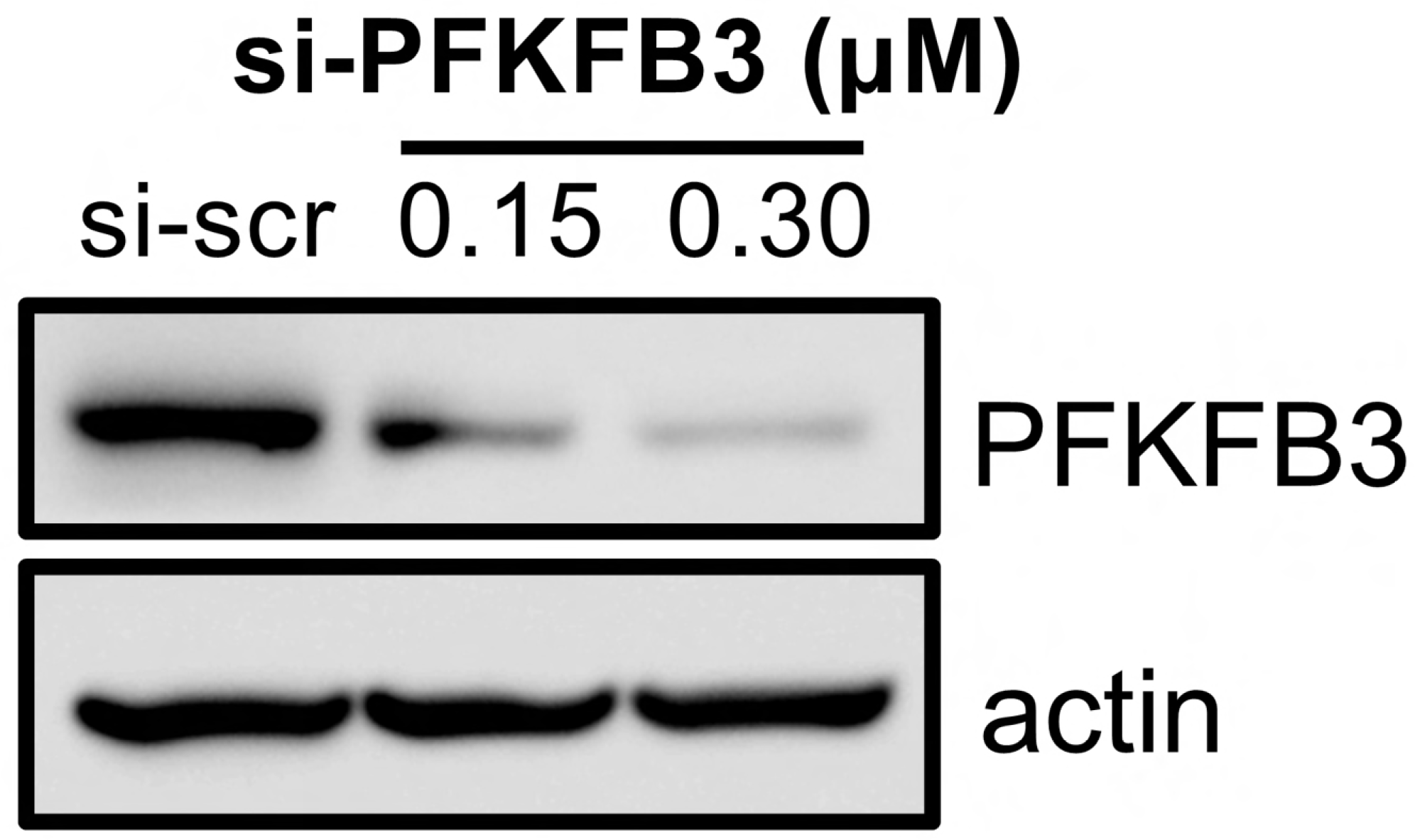
Western blot showing successful siRNA knockdown of PFKFB3 in HLMVECs. HLMVECs were transfected with PFKFB3 siRNA (0.15 and 0.3 μM) or control siRNA (0.3 μM), harvested 48 hours later for western analysis of PFKFB3 expression. Original western images are available in Figure 1 Source Data 3.

**Figure 1 Supplementary Figure 3.**
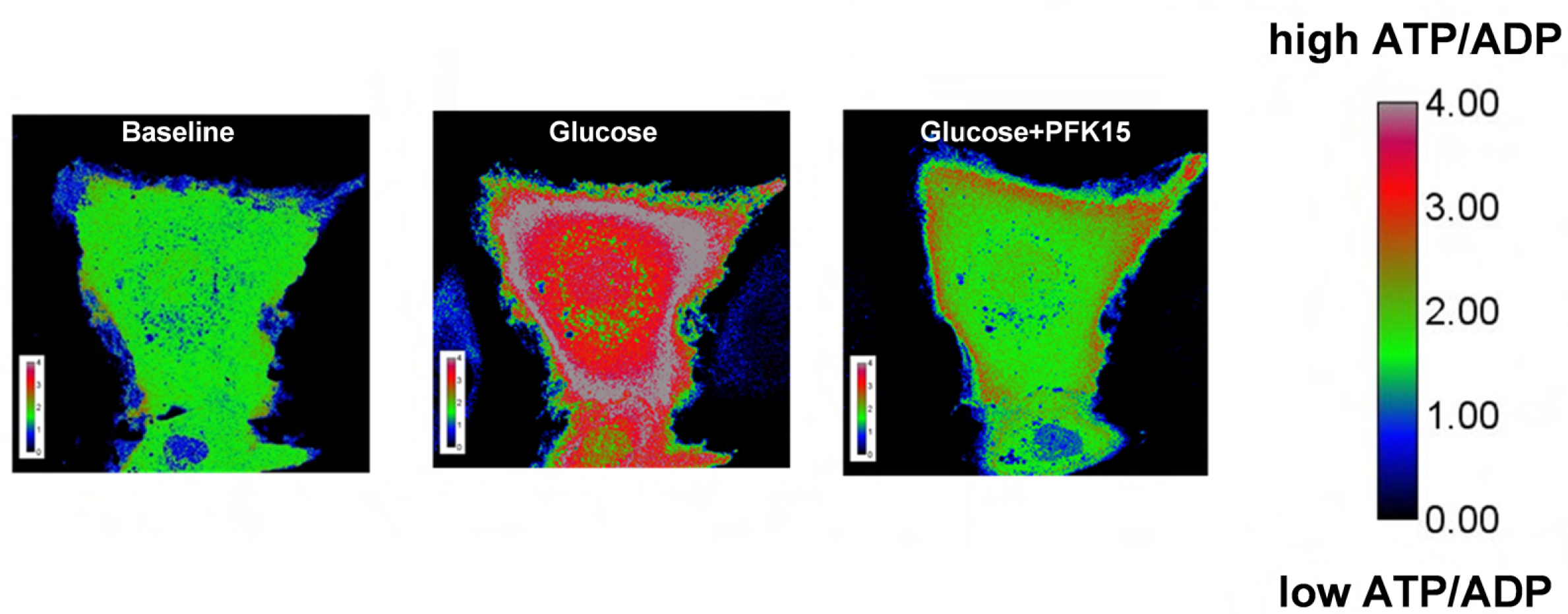
Representative images visualizing intracellular ATP/ADP levels in HLMVECs using PercevalHR. High PercevalHR F_488_/F_420_ ratio indicates high intracellular ATP/ADP. In HLMVECs expressing Perceval HR, glucose (4.5 g/L) was added first to activate glycolysis, and PFK15 (5 μM) was added later to inhibit PFKFB3. Images of glucose at 5 minutes and PFK15 after 15 minutes are shown.

**Figure 3 supplementary Figure 1.**
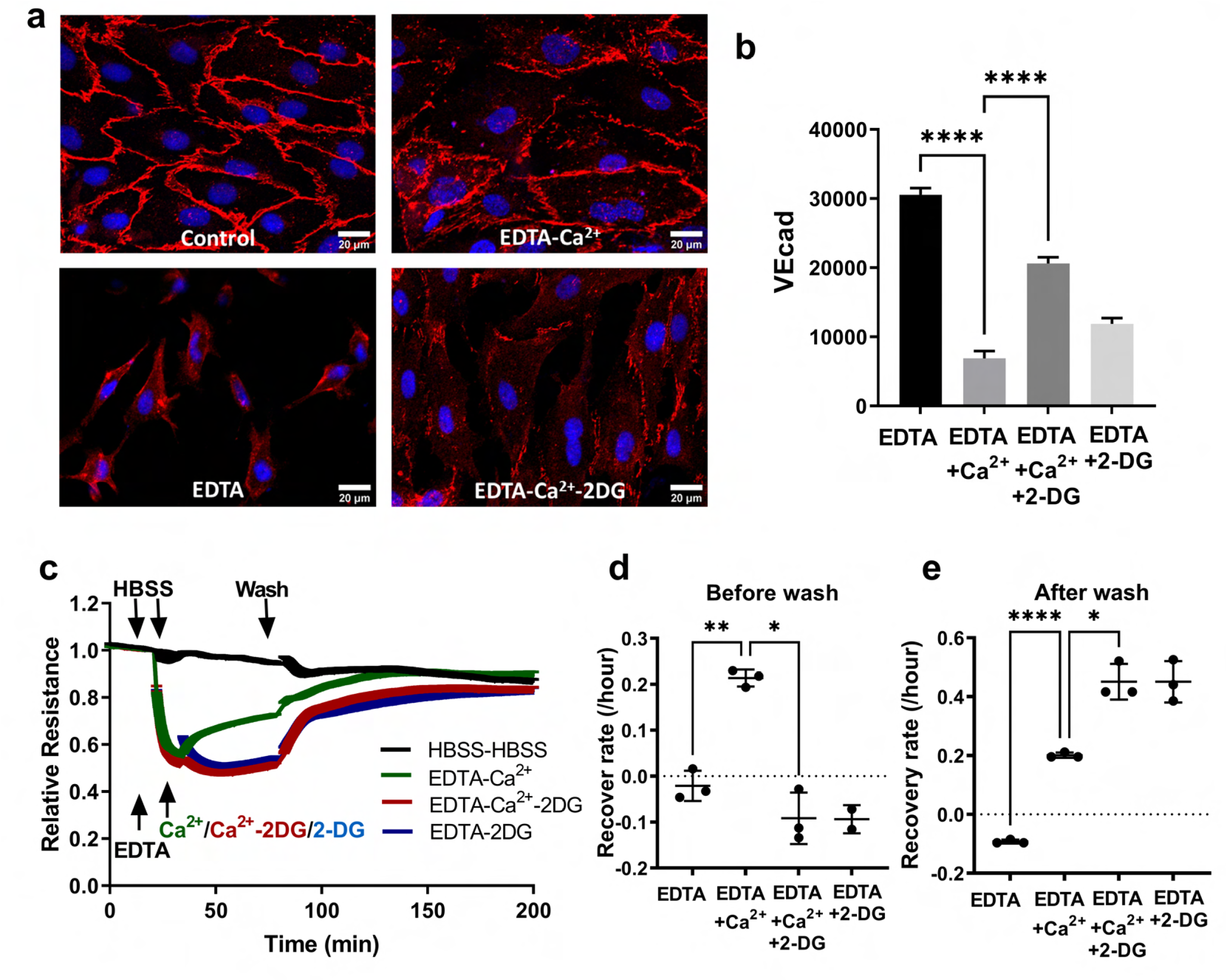
Glycolysis inhibitor 2-DG abolishes recovery of EC barrier function. (**a-b**) 2-DG prevents the re-appearance of membrane VE-cad following the addition of extracellular Ca^2+^. HLMVECs were first treated with EDTA (1mM) for 15 minutes to chelate intracellular Ca^2+,^ resulting in disruption of membrane VE-cad, then treated with Ca^2+^ or Ca^2+^/2-DG for 30 minutes. Cells were fixed and stained for membrane expression of VE-cad. Representative images are shown in a and quantification of total fluorescence of membrane VE-cad per cell is shown in b. (**c-e**) 2-DG abolishes the recovery of EC barrier function in EDTA-Ca^2+^ setting. Stabilized HLMVECs in HBSS were treated with EDTA (1mM) to disrupt the junction and reduce the resistance, 1mM Ca^2+^ to trigger the repair. 2-DG (100mM) was added together with Ca^2+^. Representative TEER measurement is shown in b and quantification of recovery rate before wash is shown in d and after wash in e. Source data for (b), (c), (d), (e), (f) and (g) are available in Figure 3 Source data 2.

**Figure 3 supplementary Figure 2.**
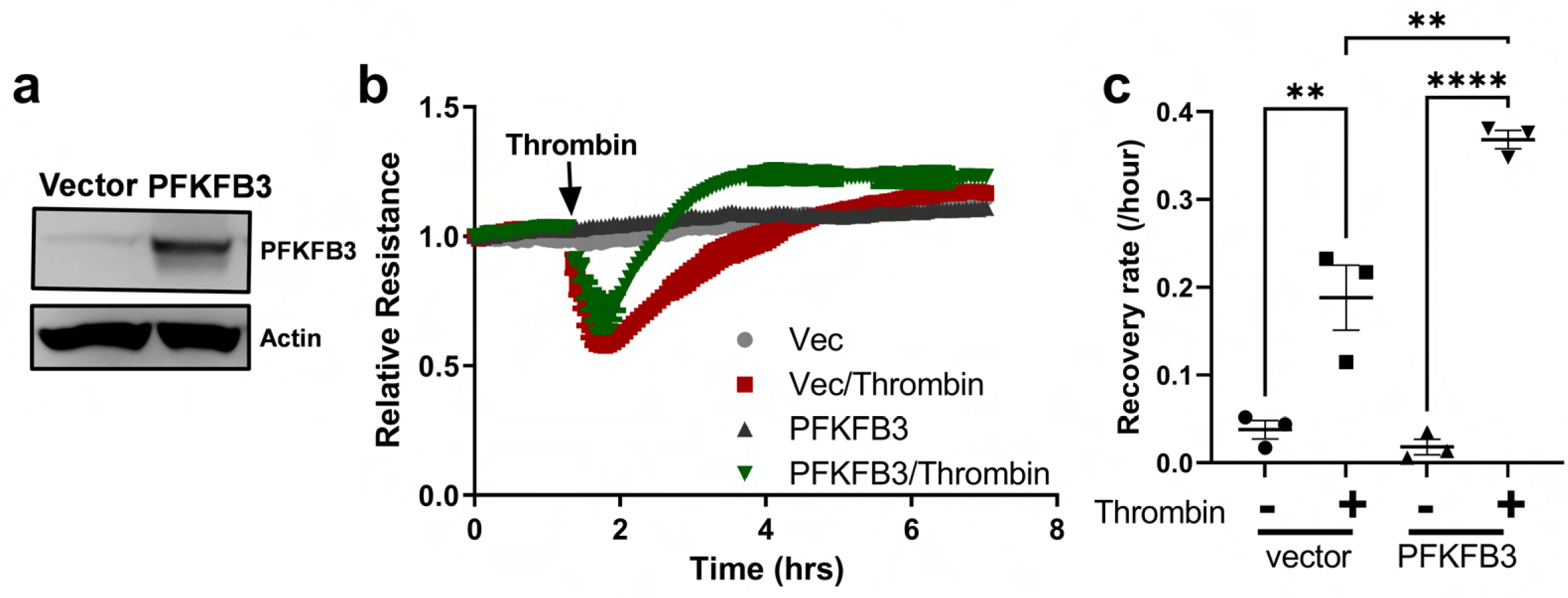
PFKFB3 enhances recovery of EC barrier function. HLMVECs were infected with lentiviral PFKFB3, using lentivirus with an empty vector as a control. 48 hours later, the cells were harvested for western blot and TEER analysis. (**a**) Western blot shows successful overexpression of PFKFB3 in HLMVECs. (**b**) Representative TEER measurement shows enhanced recovery of thrombin-induced EC permeability. HLMVECs in TEER plates were treated with Thrombin at 1unit/ml. (**c**) Quantification of TEER recovery rate from three independent experiments. Raw western blot images for (a) are available in Figure 3 Source data 3. Source data for (b) and (c) are available in Figure 3 Source data 4.

**Figure 4 supplementary Figure 1.**
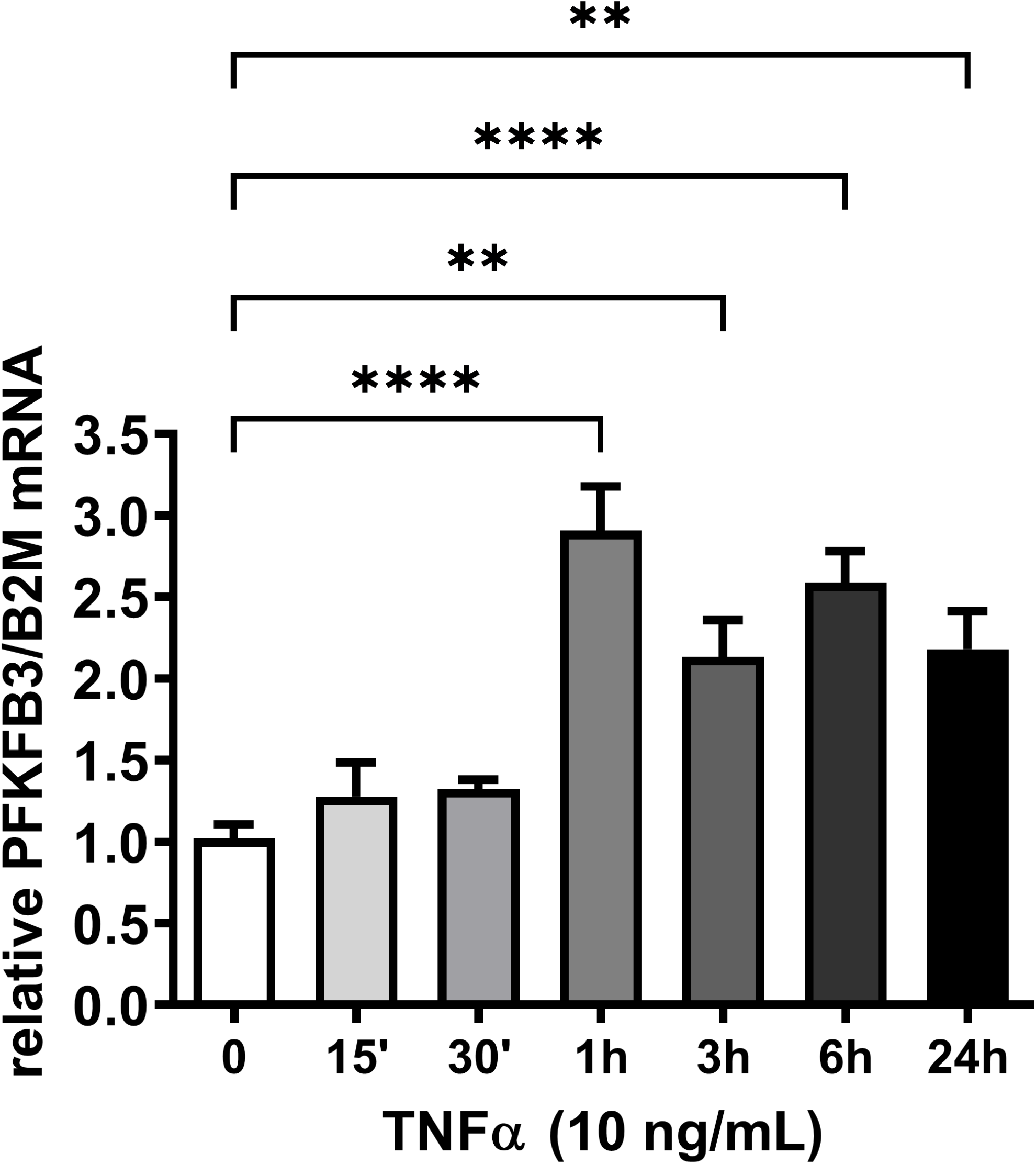
TNFα increases PFKFB3 mRNA expression. HLMVECs were treated with TNFα, at indicated time, cells were harvested for mRNA isolation, cDNA conversion, and qPCR. Source data is available in Figure 4 Source data 3.

**Figure 4 supplementary Figure 2: Western blot shows successful Hif1α knockdown.** HLMVECs were transfected with specific Hif1α siRNAs for knockdown, using non-specific siRNA as a negative control. 48 hours later, transfected cells were treated with DMOG (1 mM), using DMSO as a vehicle control, for 6 hours and harvested for western blot analysis. Raw western blot image is available in Figure 4 Source data 4.

**Figure 5 supplementary Figure 1.**
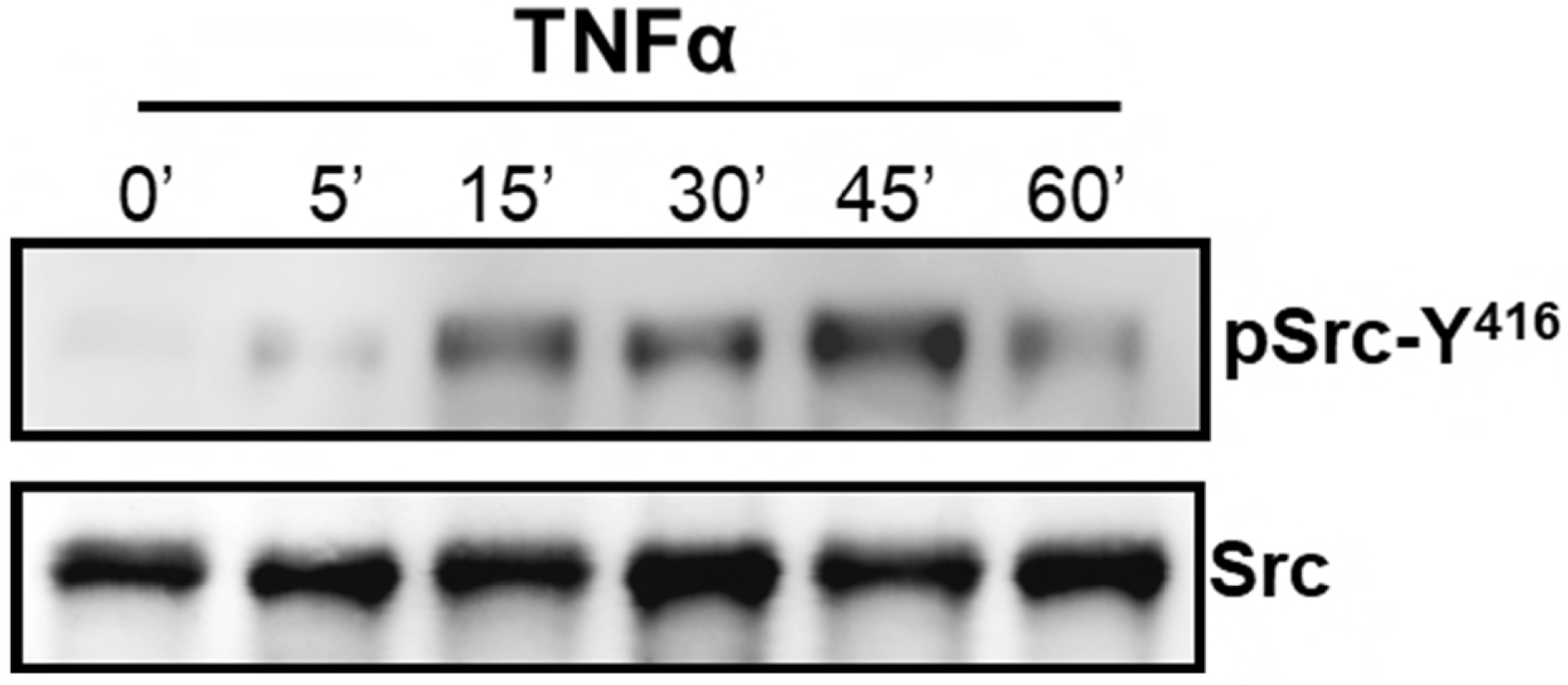
TNFα activates Src in HLMVECs. Representative western blot images showing that TNFα activates Src. Tyrosine phosphorylation at 416 (Y^416^) was used as an indicator for Src activation. Raw western blot images are available in Figure 5 Source data 3.

**Figure 5 supplementary Figure 2.**
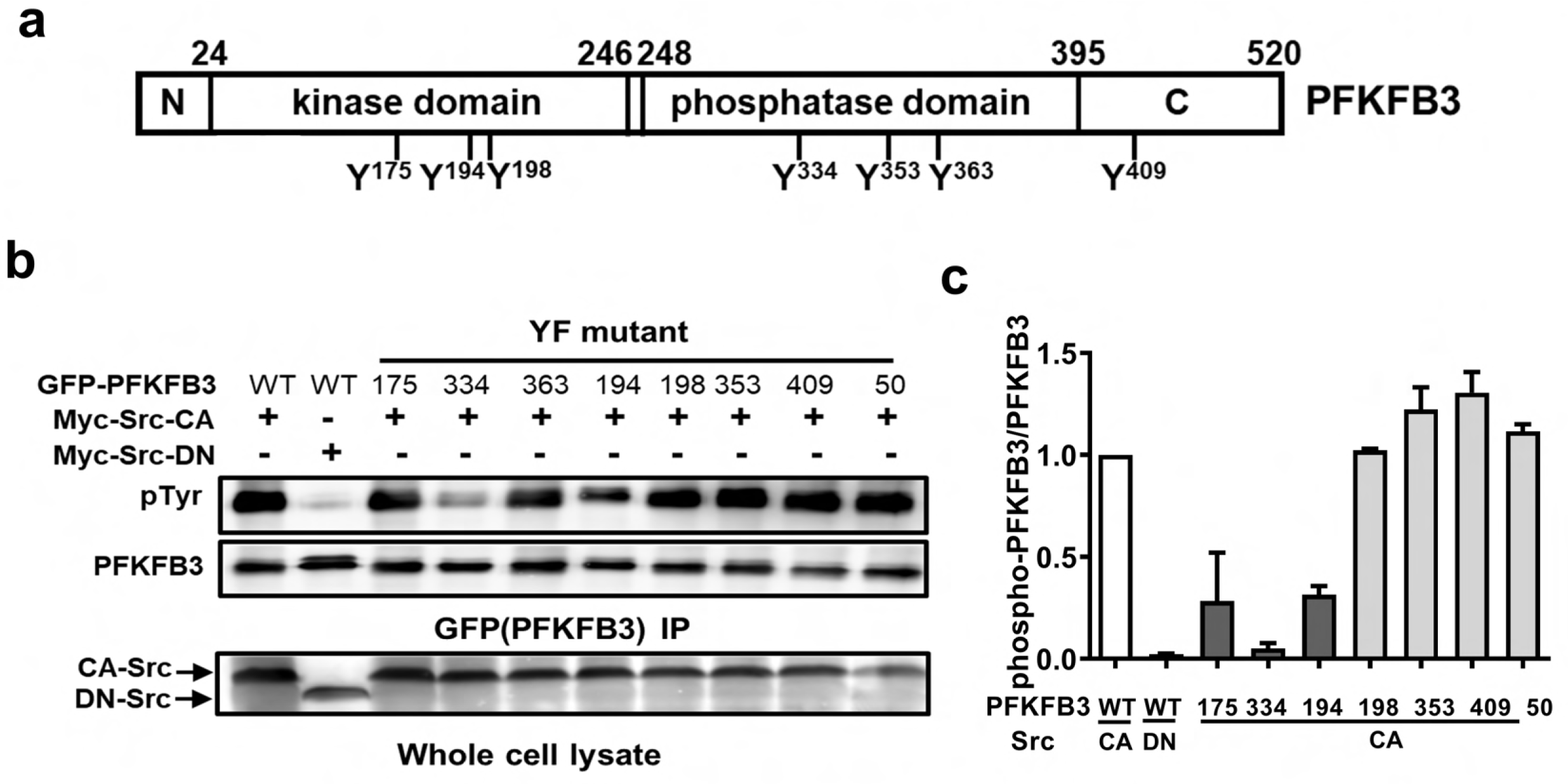
Identification of PFKFB3 tyrosine residues phosphorylated by Src. (**a**) Schematic of PFKFB3 domains and its potential tyrosine residues phosphorylated by Src. It is divided into N-terminal regulatory domain, kinase domain, phosphatase domain and C-terminal regulatory domain. On-line open access server NetPhos 2.0 was used to predict potential Src substrates. 7 residues were identified and listed, as well as Y50, which was not identified as Src substrate and was used as a negative control. (**b**,**c**) Co-expression of CA-Src with GFP-PFKFB3 showing that Src phosphorylates tyrosine at 175, 334, 363, and 194. HEK293T was co-transfected with Src (either dominant-negative or constitutively active form) and GFP-PFKFB3 (WT or mutants Y175F, Y334F, Y363F, Y194F, Y198F, Y353F, Y409F, Y50F) for expression. Cell lysate from the transfected cells were IP-ed with GFP antibody and its phosphorylation was studied using 4G10. The successful expression of Src in the whole cell lysate is also shown. Representative western blot images are shown in b. The blot was quantified using ImageJ and the relative level of tyrosine-phospho-PFKFB3 over PFKFB3 was plotted and shown in c. Raw western blot images for (b) are available in Figure 5 Source data 4.And source data for (c) is available in Figure 5 Source Data 5.

**Figure 6 supplementary Figure 1.**
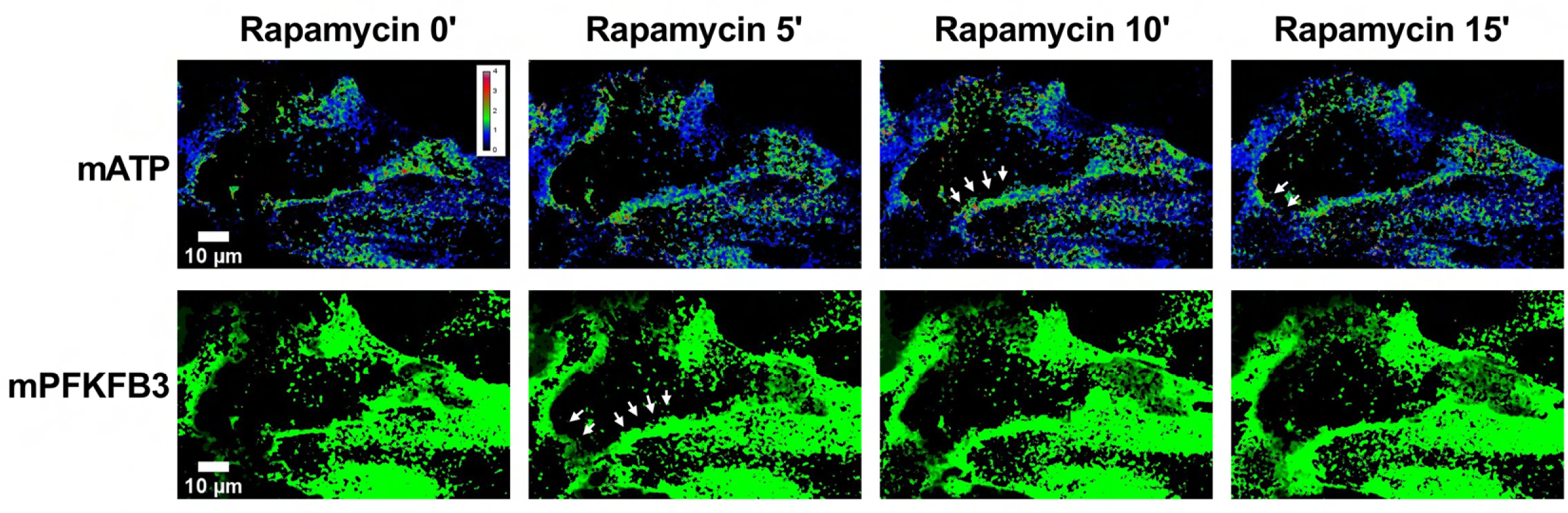
Representative images of RapT showing that rapamycin induces PFKFB3 translocation to plasma membrane and increases plasma membrane ATP. HLMVECs were infection of stargazin-iRFP-FRB and nuc-free PFKFB3-mApple-FKBP12 for expression, and images were taken every 5 minutes before and after rapamycin treatment. The 1^st^ row shows calculated mATP, the 2^nd^ row shows calculated mPFKFB3 images. The white arrows show the ATP levels at sites of cell-cell interaction.

**Figure 6 supplementary Figure 2.**
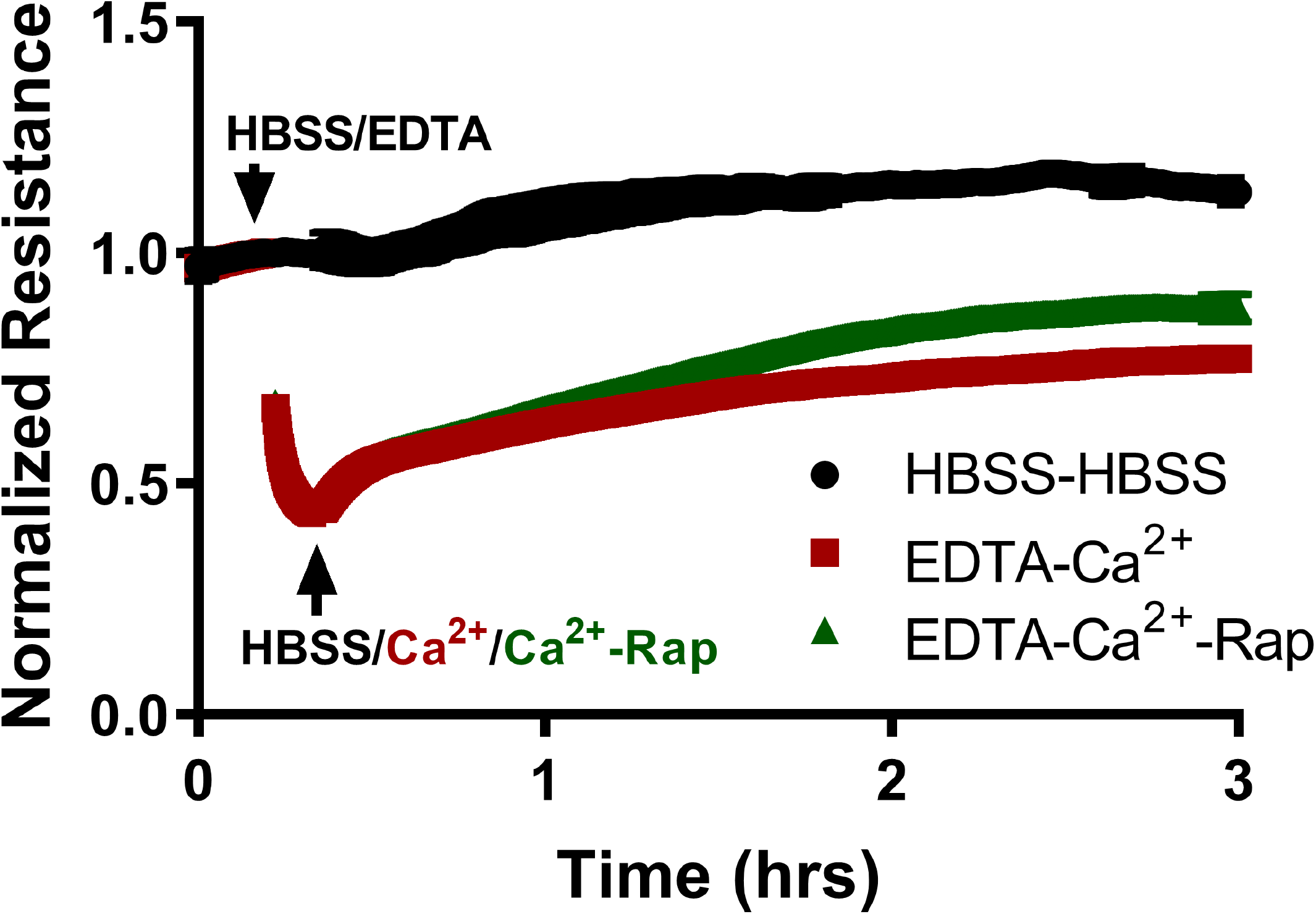
Representative TEER showing membrane PFKFB3 enhancing the recovery of EC barrier function. Cells were treated with EDTA to induce permeability, Rapamycin (0.5 μM) or Ethanol (0.05%, vehicle control) was added simultaneously with Ca^2+^ to study the effect of membrane PFKFB3 on the recovery of EB barrier function. Source data is available in Figure 6-Source data 2.

## Source Data legends Figure 1-Source Data 1

Excel files containing the original source data for figure 1a. b. c, e, g.

**Figure 1-Source Data 2**

Excel file containing the original source data for Figure 1 supplementary figure 1.

**Figure 1-Source Data 3**

Uncropped western blot images for Figure 1 supplementary figure 2.

**Figure 2 – Source Data 1**

Excel files containing the original source data for Figure 2a. b. d, g.

**Figure 3 – Source Data 1**

Excel files containing the original source data for in Figure 3b. c. d, e, f, g.

**Figure 3 - Source Data 2**

Excel files containing the original source data for Figure 3 supplementary 1b. c. d, e.

**Figure 3 - Source Data 3**

Uncropped western blot images for Figure 3 supplementary figure 2a.

**Figure 3 - Source Data 4**

Excel files containing the original source data for Figure 3 supplementary 2b, c.

**Figure 4 - Source Data 1**

Uncropped western blot images for Figure 4a, b.

**Figure 4 - Source Data 2**

Excel files containing the original source data for Figure 4a, b, c, d c.

**Figure 4 – Source Data 3**

Excel file containing the original source data for Figure 4 supplementary figure 1.

**Figure 4 – Source Data 4**

Uncropped western blot images for Figure 4 supplementary figure 2.

**Figure 5 – Source Data 1**

Uncropped western blot images for Figure 5b, d, e, f, g.

**Figure 5 – Source Data 2**

Excel files containing the original source data for Figure 5c, h, i, j.

**Figure 5 – Source Data 3**

Uncropped western blot images for Figure 5 supplementary figure 1.

**Figure 5 – Source Data 4**

Uncropped western blot images for Figure 5 supplementary figure 2b.

**Figure 5 – Source Data 5**

Excel file containing the original source data for Figure 5 supplementary figure 2c.

**Figure 6 – Source Data 1**

Excel files containing the original source data for Figure 6b, e, f, g, h, i.

**Figure 6 – Source Data 2**

Excel file containing the original source data for Figure 6 supplementary data 2.

